# Inhibition of HVEM suppresses growth and invasion of mesenchymal glioblastoma

**DOI:** 10.64898/2026.01.16.699674

**Authors:** Ryo Tanabe, Bengt Westermark, Carl-Henrik Heldin, Kohei Miyazono

## Abstract

Mesenchymal glioblastoma is a subtype of glioblastoma multiforme (GBM) characterized by pronounced inflammatory features and resistance to conventional therapies. Proneural GBM acquires a mesenchymal phenotype through proneural-mesenchymal transition (PMT), in which NF-κB signaling plays a central role. Through RNA-sequencing analysis of glioma-initiating cells (GICs), we found that expression of herpes virus entry mediator (HVEM or TNFRSF14) is highly expressed in mesenchymal GBM cells. Functional analyses revealed that HVEM promotes GIC proliferation, neurosphere formation, and invasive capacity *in vitro*, and enhances tumor formation following intracranial transplantation of GICs in mice. Among the TNF superfamily ligands, <u>a</u> <u>pr</u>oliferation-inducing ligand (APRIL or TNFSF13) binds to HVEM and activates NF-κB signaling, thereby inducing a mesenchymal phenotype in GBM cells. Furthermore, HVEM expression contributed to resistance to anticancer drugs, which was relieved by knockout of HVEM expression in the mesenchymal GICs. To therapeutically target this pathway, we generated nanobodies from camelid-derived heavy-chain-only antibodies against human HVEM. An anti-human HVEM nanobody, which binds to the cysteine-rich domain 1 (CRD1) of human HVEM, significantly inhibited the invasion of mesenchymal GICs in organotypic cultures and suppressed tumor growth in a mouse xenograft model. In addition to APRIL, HVEM binds to multiple ligands, of which B and T lymphocyte attenuator (BTLA) plays a critical role in immune evasion via binding to HVEM. The anti-human HVEM nanobody blocked interaction between HVEM and BTLA. Collectively, these findings suggest that the anti-human HVEM nanobody regulates multiple signaling pathways, and that HVEM represents a promising therapeutic target for the treatment of mesenchymal GBM.

**One Sentence Summary:** HVEM drives aggressive glioblastoma by boosting tumor growth and invasion upon binding of APRIL, while an anti-HVEM nanobody slows tumor progression.

## INTRODUCTION

Glioblastoma multiforme (GBM) is the most common and malignant form of brain tumors in adults. GBM cells proliferate rapidly, infiltrate extensively, and typically show marked nuclear pleomorphism and necrosis. Despite advances in surgery, radiotherapy, and chemotherapy, as well as new treatment strategies, the median survival of patients with GBM remains less than 20 months (*1, 2*). Recent efforts to target abnormalities in the signaling pathways in GBM cells have not been successful in improving treatment efficacy (*3–5*).

Glioma-initiating cells (GICs) constitute a minor subpopulation of GBM cells, playing central roles in the initiation, progression, recurrence, tumor heterogeneity, and drug and radiation resistance of GBM (*6, 7*). GBM has been classified into subtypes defined by transcriptomic profiles, such as proneural (PN), classical (CL), and mesenchymal (MES) profiles (*8, 9*). The originally reported neural subtype (NL) (*8*) lacks a distinct gene signature and is currently regarded as a non-tumor-specific phenotype, reflecting contamination of increased normal neural tissues at the tumor margin (*8*). Single-cell RNA sequencing has demonstrated that GBM cells exist in four main cellular states: neural progenitor-like (NPC-like), oligodendrocyte progenitor-like (OPC-like), astrocyte-like (AC-like), and mesenchymal-like (MES-like) (*10*). The features of non-MES subtypes are linked to genetic driver genes of GBM, such as *CDK4*, *PDGFRA*, and *EGFR*, whereas the mesenchymal subtype often involves *NF1* alterations (*8, 10*). Although classifications of GBM tissues and cells based on their molecular profiles are widely used, the pronounced plasticity of GBM cells allow cells of different subtypes to co-exist in the same tumor (*11, 12*). Moreover, shift in subtype can occur in patients with GBM (*13*, *14*).

Proneural GBM cells often undergo transition to mesenchymal states characterized by enhanced inflammatory features through proneural-mesenchymal transition (PMT) (*15*-*17*). Mesenchymal transition in GBM has been reported to be promoted by certain transcription factors, such as STAT3, CEBPβ, TAZ, NF-κB, c-FOS, and c-JUN (*13, 18*-*21*). The mesenchymal profile of GBM is associated with high invasiveness and resistance to drug and radiation therapy (*16*). Thus, strategies to treat the mesenchymal subtype of GBM are required to improve the survival of GBM patients.

Interactions between GBM cells and the tumor microenvironment are critical determinants of the mesenchymal phenotype. Among various cells in the tumor microenvironment, macrophages appear to be key inducers of the mesenchymal-like state in GBM cells (*10*, *21*). Notably, *NF1* deficiency observed in mesenchymal GBM cells promotes the recruitment of tumor-associated macrophages into the microenvironment, and is associated with increased resistance to radiotherapy (*9*). Among the transcription factors which have been suggested to be involved in mesenchymal transition, NF-κB is of particular interest (*21*, *22*). Although several tumor necrosis factor (TNF) superfamily factors, including TNF-α and TWEAK (TNF-like weak inducer of apoptosis), may be involved in GBM progression (*13*, *22*, *23*), specific factor(s) responsible for NF-κB activation during the induction of PMT remain undefined.

Here, we show that expression of herpes virus entry mediator (HVEM, or TNF receptor superfamily member 14; TNFRSF14) is enhanced in mesenchymal GICs and contributes to progression of GBM. Intriguingly, <u>a</u> <u>pr</u>oliferation inducing ligand (APRIL, or TNF superfamily member 13; TNFSF13) is secreted by GICs and physically interacts with HVEM, leading to activation of NF-κB signaling. Thus, our findings suggest a critical role of tumor-intrinsic HVEM in progression of mesenchymal GICs through its interaction with APRIL. We generated an anti-human HVEM nanobody, which significantly inhibited the invasion and tumor growth of mesenchymal GICs. Among multiple ligands binding to HVEM, B and T lymphocyte attenuator (BTLA) is critically involved in immune evasion. In addition to its effects on growth and invasion of mesenchymal GICs, we found that the anti-human HVEM nanobody blocked interaction of HVEM with BTLA.

## RESULTS

### *HVEM*/*TNFRSF14* is highly expressed in the mesenchymal GBM cells

To identify molecular targets in mesenchymal GBM for antibody-based therapies, genes encoding membrane proteins that are expressed differentially in mesenchymal and non-mesenchymal GBM were extracted from the dataset of the human glioblastoma cell culture (HGCC) Resource (*24, 25*), and their expression levels were compared between non-mesenchymal and mesenchymal cells (Fig. 1A). Since PMT in GBM cells is induced by NF-κB signaling (*13, 16*), we were interested in molecule(s) which are highly expressed in mesenchymal but not in non-mesenchymal GICs, and can activate NF-κB signaling. *HVEM*/*TNFRSF14* encodes a member of the TNF receptor superfamily, that is well known to induce NF-κB signaling upon ligand binding (*26*, *27*). The expression of *HVEM/TNFRSF14* was determined in 13 GICs from the HGCC Resource; *HVEM* was found to be highly expressed in mesenchymal GICs compared to non-mesenchymal GICs (Fig. 1B). Consistently, analysis of The Cancer Genome Atlas (TCGA) dataset demonstrated that *HVEM*/*TNFRSF14* was distinctly expressed in mesenchymal GBM tissues (Fig. 1C). Lack of *IDH1/2* mutations and low expression of glioma CpG island methylator phenotype (G-CIMP) are known to correlate with high malignancy of brain tumors (*28*-*30*). We found that expression levels of *HVEM*/*TNFRSF14* correlated with lack of *IDH* mutations and low expression of G-CIMP (Fig. 1D and 1E). Moreover, high expression of *HVEM*/*TNFRSF14* correlated with poor prognosis of GBM patients (Fig. 1F). These findings suggested that targeting of HVEM may be beneficial in the treatment of patients with mesenchymal GBM.

**Fig. 1.**
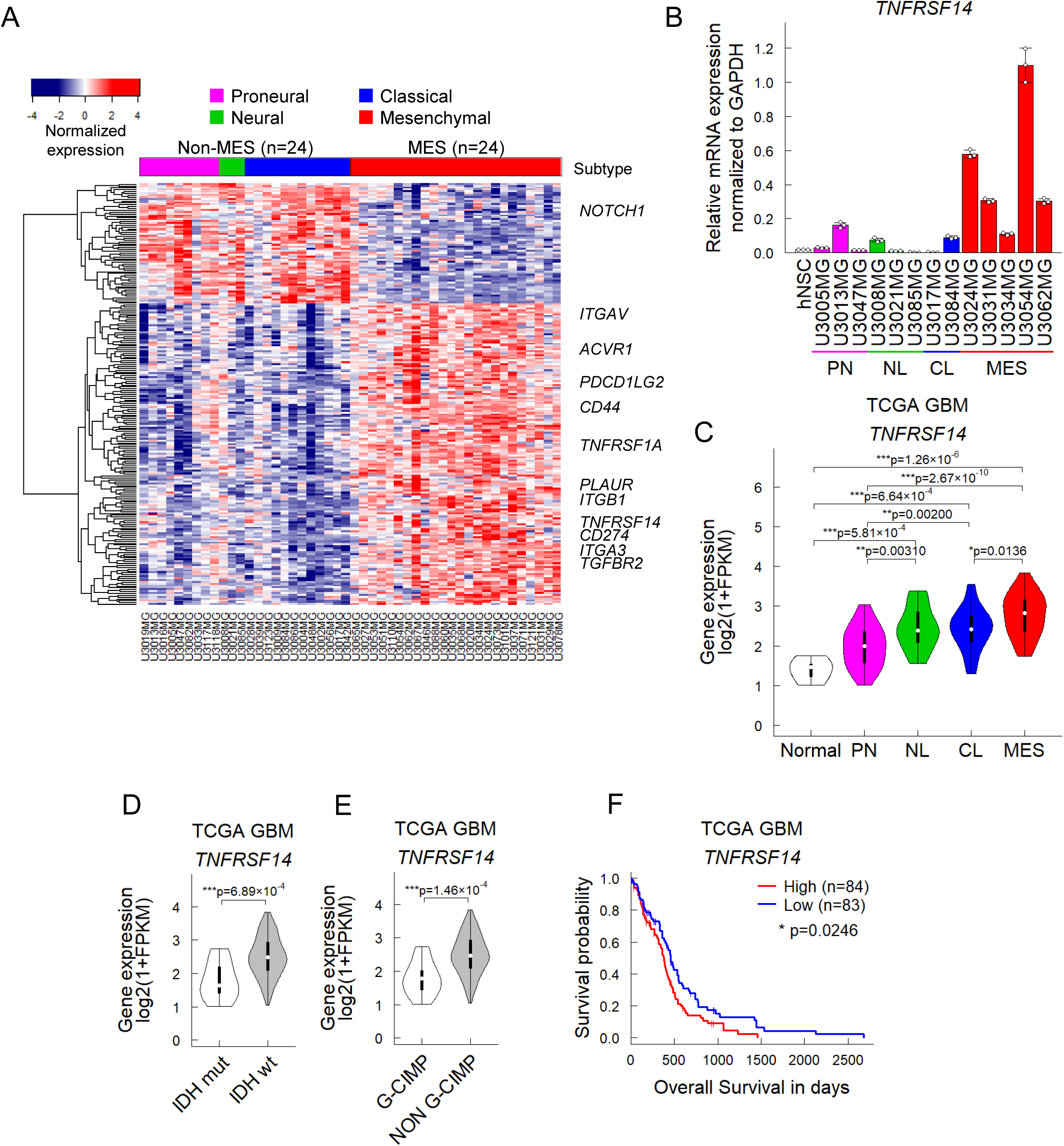
*HVEM/TNFRSF14* expression levels are elevated in malignant brain tumors and correlate with poor prognosis. (**A**) A heatmap of differentially expressed genes encoding membrane proteins between mesenchymal and non-mesenchymal GBM from GSE72217. (**B**) Expression levels of *TNFRSF14* among human neural stem cells (hNSCs) and 4 subtypes of human glioblastoma-initiating cells (GICs). GBM cell lines in the HGCC Resource were analyzed for the expression of *TNFRSF14* by quantitative real-time PCR. Data are shown as mean ± SD (n=3 biological replicates). PN, proneural; NL, neural; CL, classical; MES, mesenchymal. (**C**) Expression levels of *TNFRSF14* in normal brain and brain tumor tissues in The Cancer Genome Atlas (TCGA) dataset (\**P*<0.05, \*\**P*<0.01, \*\*\**P*<0.001; two-tailed Kruskal-Wallis test with Bonferroni’s correction). (**D, E**) Expression levels of *TNFRSF14* in subtypes of GBM tissues in the TCGA dataset (\*\*\**P*<0.001; two-tailed Kruskal-Wallis test). Brain tumor tissues with or without *IDH* mutations (**D**) and those with or without glioma CpG island methylator phenotype (G-CIMP) (**E**) were analyzed. (**F**) Kaplan-Meier plot of the survival of GBM patients in TCGA dataset. The patients were equally divided into two groups based on *TNFRSF14* expression level (\**P*<0.05; two-tailed log-rank test).

Bone morphogenetic proteins (BMPs) induce differentiation, cell cycle arrest, and apoptosis of GICs (*31*-*33*). However, a subset of GICs, particularly mesenchymal GICs, acquire resistance to BMP signaling (*34*). We found that BMP-4 inhibited proliferation and sphere formation of mesenchymal U3024MG and U3031MG cells, whereas no such effects were observed in U3054MG cells (fig. S1A, B). Consistent with these findings, BMP-4 suppressed the expression levels of *HVEM* in BMP-responsive cells (fig. S1C), suggesting that HVEM is involved in the promotion of pro-tumorigenic properties, including stem cell-like characteristics, in GICs.

### Disruption of *HVEM* expression attenuates tumor progression of mesenchymal GICs

To investigate the function of HVEM, the *HVEM/TNFRSF14* gene was knocked down by short hairpin RNA (shRNA), or edited by CRISPR/Cas9 technology, in mesenchymal GICs. Decreased expression of HVEM on the cell surface of mesenchymal GICs after knockdown by shRNA, or knockout by single guide RNA (sgRNA), was confirmed by flowcytometric analysis (Fig. 2A). Knockdown or knockout of *HVEM* inhibited *in vitro* cell proliferation of mesenchymal GICs (Fig. 2B), and decreased the number of proliferating cells in a 5-ethynyl-2’-deoxyuridine (EdU)-incorporation assay (Fig. 2C). Moreover, sphere formation of mesenchymal GICs was suppressed by knockdown or knockout of *HVEM*, suggesting that stem cell frequency was decreased by loss of the HVEM expression (Fig. 2D).

**Fig. 2.**
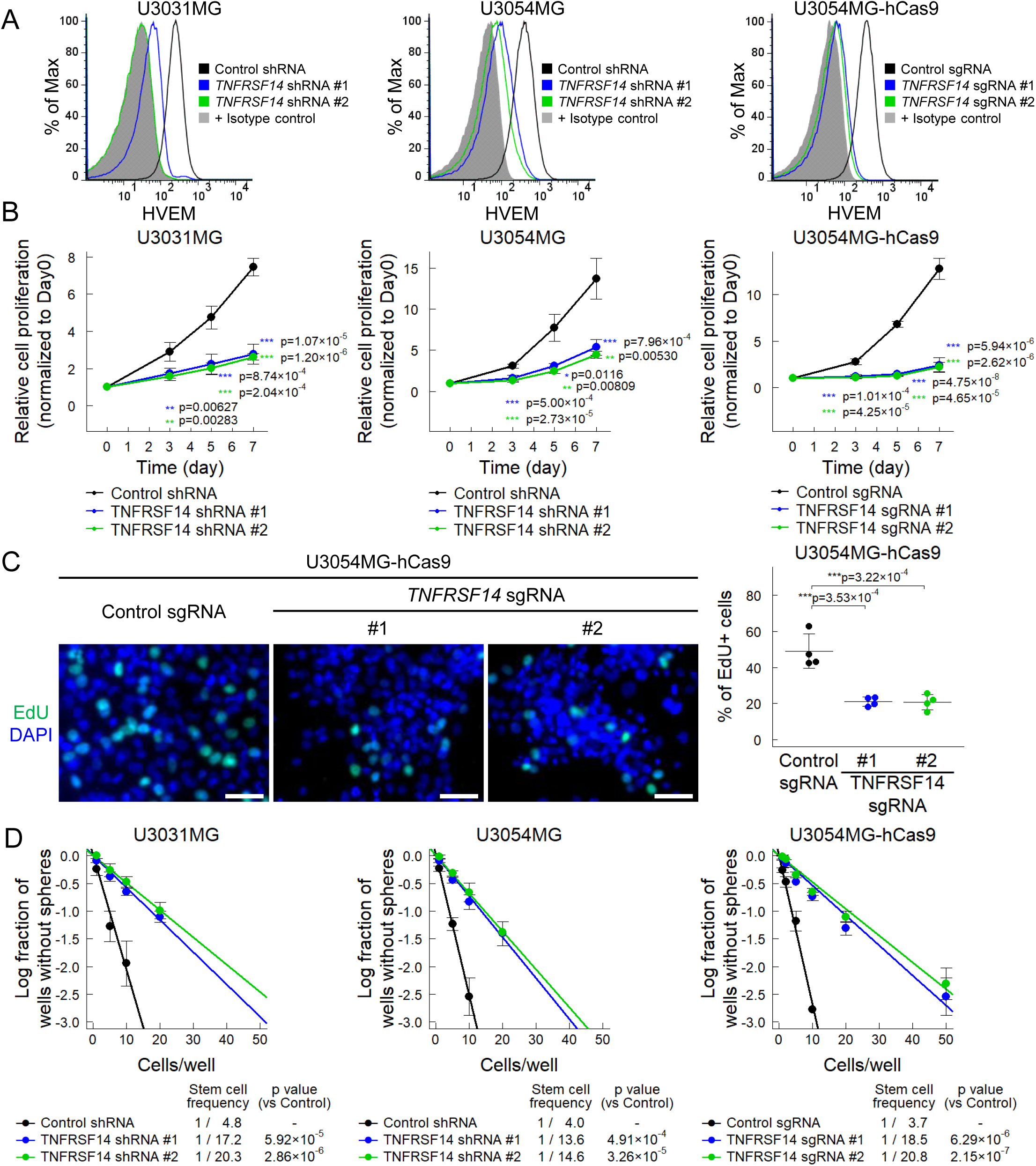
Genetic disruption of *HVEM/TNFRSF14* attenuates the GIC phenotype. (**A**) Cell surface expression of HVEM in mesenchymal GBM cells upon genetic disruption. The *HVEM/TNFRSF14* gene was knocked down in mesenchymal GICs (U3031MG and U3054MG) by lentivirus-mediated shRNA, or edited in U3054MG cells by lentivirus-mediated CRISPR-Cas9 system (U3054MG-hCas9). Cell surface expression of HVEM was evaluated by flowcytometric analysis. (**B**) Growth curves of mesenchymal GICs expressing shRNA or combination of Cas9 and sgRNA. Data are shown as mean ± SD (n=4 biological replicates; \*\**P*<0.01, \*\*\**P*<0.001; Tukey’s HSD test). (**C**) Analysis of proliferative ability by EdU labeling of U3054MG cells. The incorporated EdU was visualized by using EdU-Click 647 kit (green) and nuclei were stained with DAPI (blue). Scale bar: 50 μm. (left). Quantification data of EdU-positive cells are shown as mean ± SD (n=4 biological replicates; \*\*\**P*<0.001; Tukey’s HSD test) (*right*). (**D**) Sphere formation ability of mesenchymal GICs expressing *HVEM/TNFRSF14* shRNA or combination of Cas9 and sgRNA. Sphere formation of mesenchymal GICs was evaluated by limiting dilution assay. Data are shown as mean ± SD (n=3 independent experiments; \*\**P*<0.01, \*\*\**P*<0.001; two-way ANOVA with Bonferroni’s correction).

To determine the effect of HVEM depletion on tumor progression *in vivo*, mesenchymal GICs in which *HVEM* had been knocked down by shRNAs were injected intracranially into nude mice. *In vivo* bioluminescent imaging demonstrated that shRNA-mediated knockdown of *HVEM* decreased the tumorigenic activity of two mesenchymal GICs, i.e., U3031MG and U3054MG cells (Fig. 3A). Furthermore, knockdown of *HVEM* in the mesenchymal GICs prolonged the survival of mice bearing tumors (Fig. 3B).

**Fig. 3.**
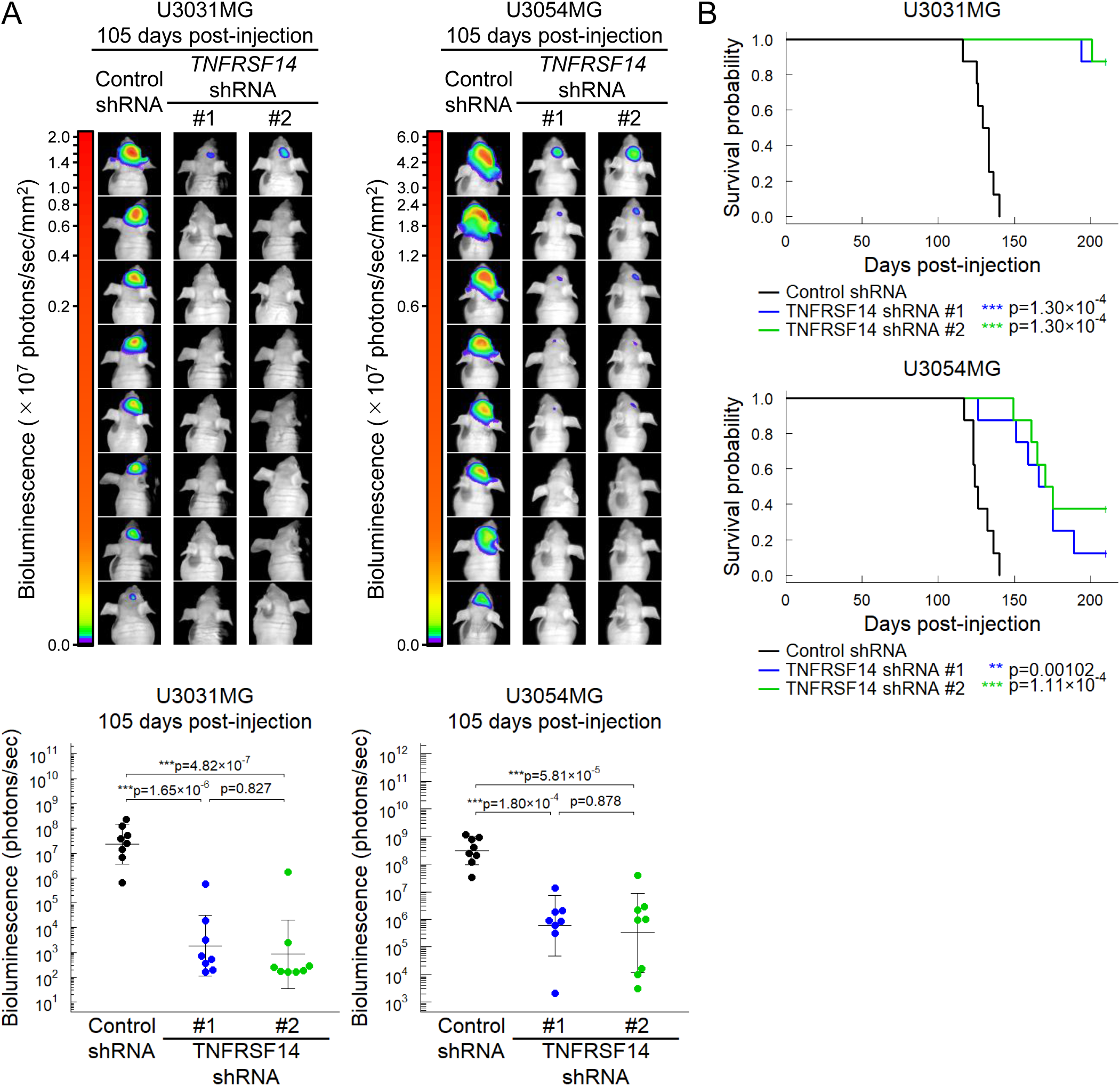
Genetic silencing of *HVEM/TNFRSF14* attenuates *in vivo* tumorigenic activity of mesenchymal GICs. (**A**) *In vivo* bioluminescent imaging of mesenchymal GICs in which *TNFRSF14* had been knocked down by shRNA, using the *firefly* luciferase method. Quantification data of bioluminescence are shown in the lower panels, as mean ± SD (n=8 mice per group; \*\*\**P*<0.001; Tukey’s HSD test). (**B**) Survival curves of mice bearing tumors derived from shRNA-expressing mesenchymal GICs (n=8 mice per group; \*\**P*<0.01, \*\*\**P*<0.001; two-tailed log-rank test with Bonferroni’s correction).

### Ectopic expression of HVEM accelerates tumor progression of non-mesenchymal GICs

We next investigated the effect of overexpression of HVEM in GIC cells. The *HVEM* gene was transduced lentivirally into non-mesenchymal GICs; increased expression of HVEM on the cell surface was confirmed by flowcytometric analysis (Fig. 4A). Ectopic HVEM expression promoted cell proliferation and sphere formation of non-mesenchymal GICs, i.e. proneural U3047MG and classical U3017MG cells (Fig. 4B, C). Furthermore, invasive activity of the non-mesenchymal GICs analyzed by an organotypic invasion assay was significantly increased by ectopic expression of HVEM (Fig. 4D).

**Fig. 4.**
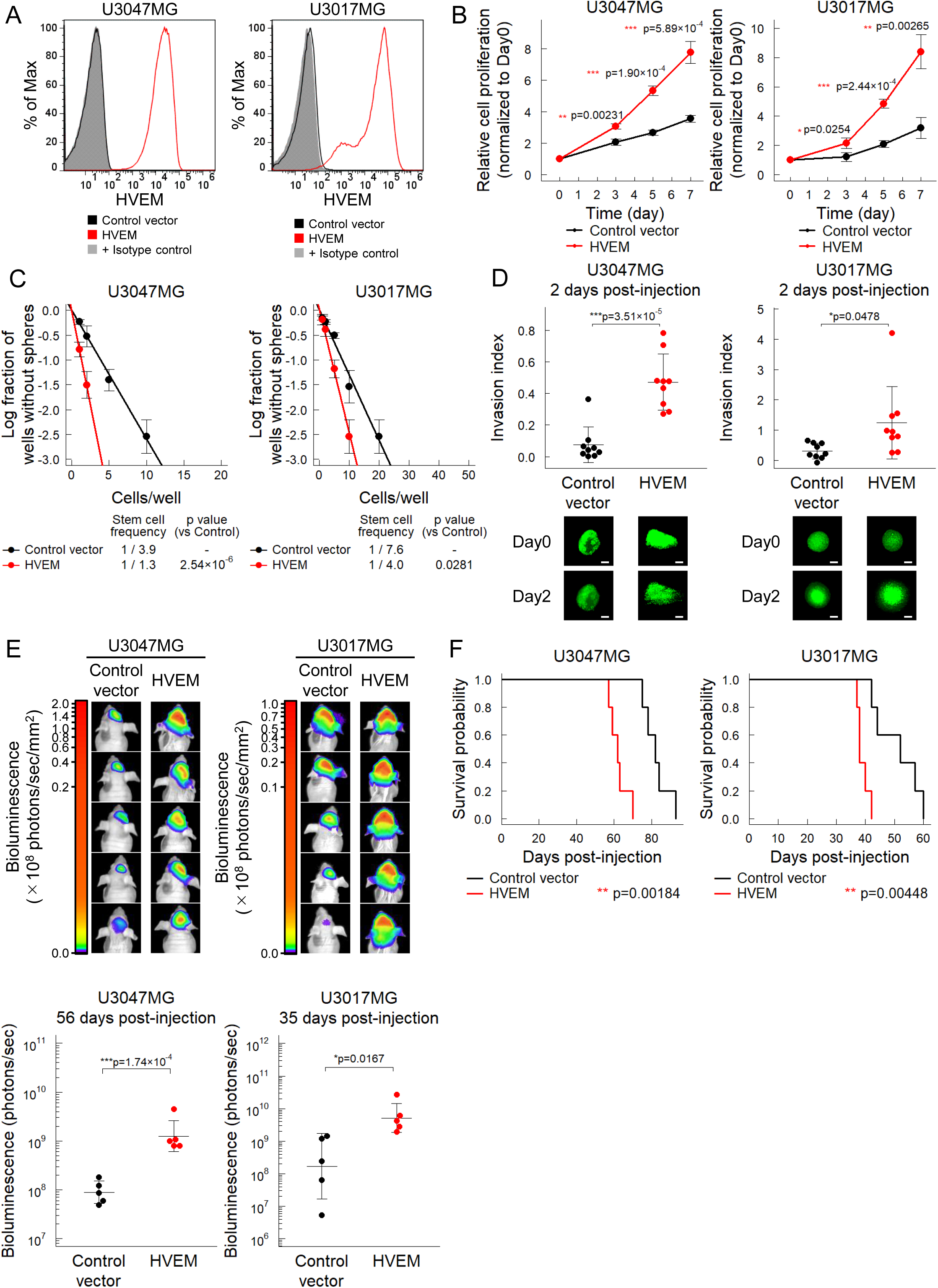
Ectopic expression of HVEM enhances tumorigenic activity of non-mesenchymal GICs. (**A**) Cell surface expression of HVEM in non-mesenchymal GICs upon ectopic expression. Ectopic HVEM was transduced lentivirally into non-mesenchymal GICs (proneural U3047MG and classical U3017MG). Cell surface expression of HVEM was evaluated by flowcytometric analysis. (**B**) Growth curves of non-mesenchymal GICs expressing HVEM or not. Data are shown as mean±SD (n=3 biological replicates; \**P*<0.05, \*\**P*<0.01, \*\*\**P*<0.001; two-tailed unpaired Student’s t-test). (**C**) Sphere forming ability of non-mesenchymal GICs expressing HVEM or not was evaluated by limiting dilution assay. Data are shown as mean ± SD (n=3 independent experiments; \**P*<0.05, \*\*\**P*<0.001; two-way ANOVA). (**D**) Invasiveness of non-mesenchymal GICs expressing HVEM was evaluated by organotypic invasion assay. Fluorescence images with median invasion index in each group are demonstrated as representative images. Quantification data are shown as mean ± SD (n=9 biological replicates; \**P*<0.05, \*\*\**P*<0.001; two-tailed unpaired Student’s t-test). Scale bar: 200 μm. (**E**) *In vivo* bioluminescent imaging of non-mesenchymal GICs expressing HVEM and *firefly* luciferase. Quantification data of bioluminescence are shown as mean ± SD (n=5 mice per group; \**P*<0.05, \*\*\**P*<0.001; two-tailed unpaired Student’s t-test). (**F**) Survival curves of mice bearing tumors derived from non-mesenchymal GICs expressing HVEM or not (n=5 mice per group; \*\**P*<0.01; two-tailed log-rank test).

To determine whether ectopic expression of HVEM affects *in vivo* tumorigenic activity, two non-mesenchymal GICs overexpressing HVEM were injected intracranially into nude mice. Ectopic HVEM expression enhanced tumor progression, as demonstrated by *in vivo* bioluminescent imaging (Fig. 4E), and shortened the survival of mice bearing tumors (Fig. 4F).

### Expression of TNF-related factors in GBM tissues and cells

TNF-related cytokines, such as LIGHT (also known as TNF superfamily 14 or TNFSF14) and lymphotoxin α (LTα, encoded by *LTA*), as well as non-TNF-related cytokines, such as BTLA, CD160, and synaptic cell adhesion molecule 5 (SALM5, encoded by *LRFN5*), are known to interact with HVEM (*26*, *27*) (Fig. 5A). To investigate which ligand(s) are responsible for the tumorigenic activities of HVEM on mesenchymal GBM cells, the expression levels of mRNAs encoding potential ligands in GBM and normal brain tissues were analyzed using the TCGA dataset (Fig. 5B). The expression of *LIGHT/TNFSF14, LTA,* and *BTLA* was significantly increased in GBM tissues, whereas that of *SALM5/LRFN5* was decreased in GBM tissues, compared to normal brain tissues. APRIL (TNFSF13) was reported previously to interact with HVEM (*35*). As shown in Fig. 5B, there was no significant difference of expression of *APRIL/TNFSF13* between GBM and normal brain tissues, but its expression levels were higher than those of the other interactants of HVEM.

**Fig. 5.**
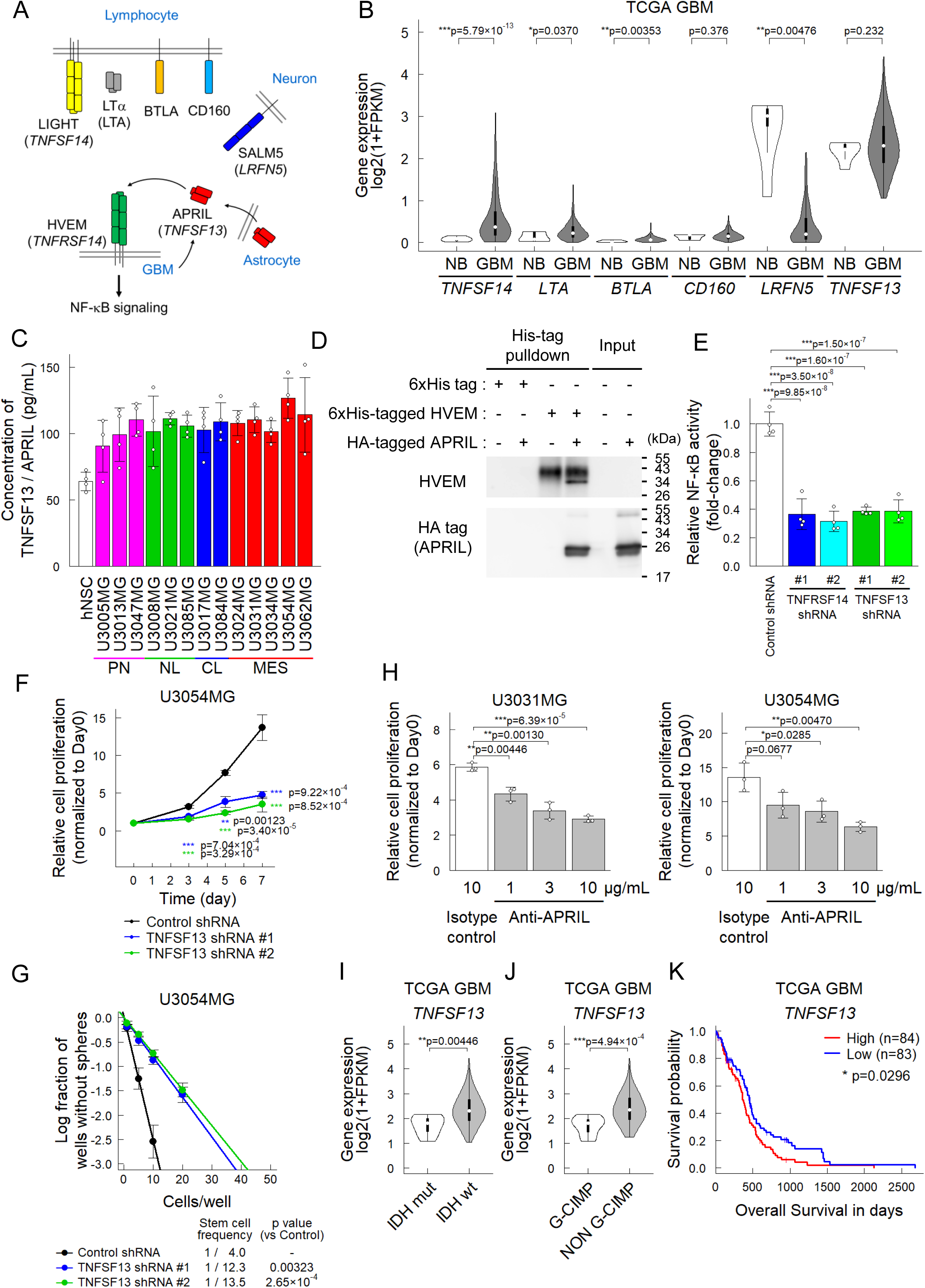
The APRIL-HVEM axis promotes progression of malignant brain tumors. (**A**) HVEM and its potential interactants. LIGHT is known to induce trimerization of HVEM, while BTLA, CD160, and SALM5 interact with a monomeric form of HVEM (*61*). (**B**) Analysis of the TCGA dataset of the expression levels of genes encoding interactants of HVEM/TNFRSF14 in brain tumor tissue. NB, normal brain (\**P*<0.05, \*\**P*<0.01, \*\*\**P*<0.001; two-tailed Kruskal-Wallis test with Bonferroni’s correction). (**C**) Analysis of APRIL concentrations in mesenchymal and non-mesenchymal GICs by sandwich ELISA. (**D**) Physical interaction between recombinant HVEM and APRIL analyzed by pulldown assay. (**E**) NF-κB activity in mesenchymal GICs upon knockdown of *HVEM*/*TNFRSF14* or *APRIL*/*TNFSF13* by shRNA (n=4 biological replicates). Data are shown as mean ± SD (\*\*\**P*<0.001; Tukey’s HSD test). (**F, G**) Growth curves (**F**; n=4 biological replicates) and sphere formation (**G**; n=3 independent experiments) of wild-type U3054MG cells and U3054MG cells expressing shRNA for *APRIL*/*TNFSF13*. Data are shown as mean ± SD (\**P*<0.05, \*\**P*<0.01, \*\*\**P*<0.001; Tukey’s HSD test (**F**) and two-way ANOVA with Bonferroni’s correction (**G**)). (**H**) Effect of a neutralizing antibody against APRIL on cell proliferation of mesenchymal GICs at day 7. Data are shown as mean ± SD (n=3 biological replicates; \*\**P*<0.01, \*\*\**P*<0.001; Tukey’s HSD test). (**I, J**) Analysis of TCGA dataset of *APRIL*/*TNFSF13* in GBM. The expression levels of *APRIL*/*TNFSF13* correlates with lack of *IDH1/2* mutations (**I**) and low G-CIMP expression (**J**) in GBM. (**K**) Kaplan-Meier plots of brain tumor patients in TCGA dataset. The patients were equally divided into two groups based on expression level of *TNFSF13* (encoding APRIL).

We also analyzed the expression of potential HVEM ligands in neural cells from humans and mice (fig. S2A). Notably, APRIL/TNFSF13 showed high expression in human mature astrocytes, but its expression was low in mouse astrocytes. In contrast, microglia/macrophages in mice exhibited relatively high expression levels of April/Tnfsf13 and Light/Tnsfsf14, whereas such expression was not observed in human cells (fig. S2A). In both species, SALM5/LRFN5 was highly expressed in neurons and oligodendrocytes.

We next examined the production of TNF superfamily ligands in human GICs using sandwich ELISA assays. APRIL was produced by hNSCs and GICs, regardless of GBM subtypes (Fig. 5C). In contrast, the secretion levels of LIGHT and LTα in GICs were lower than the limit of detection (fig. S2B).

### APRIL interacts with HVEM to induce NF-κB signaling in GBM

We investigated the functional interaction between HVEM and APRIL. Pulldown assay demonstrated physical interaction between recombinant HVEM and APRIL (Fig. 5D). To investigate whether APRIL stimulation induced intracellular signaling, NF-κB activity was measured by a reporter assay. Knockdown of *APRIL/TNFSF13* by shRNA decreased NF-κB activity as did knockdown of *HVEM/TNFRSF14* (Fig. 5E). To validate that APRIL activated NF-κB signaling through HVEM, we established HEK293T cells (target cells) expressing HVEM and an NF-κB reporter. These target cells were cocultured with HEK293T cells expressing a ligand (effector cells), including APRIL, LIGHT or SALM5 (fig. S3A). APRIL is produced exclusively as a soluble form (*36*). Interestingly, soluble APRIL elicited NF-κB activation to an extent comparable to that induced by either the soluble or membrane-bound forms of LIGHT or SALM5 (fig. S3B, C).

Similar to suppression of *HVEM,* silencing of *APRIL* by shRNA inhibited cell proliferation and sphere formation of mesenchymal GIC cells (Fig. 5F and 5G). Consistently, *in vitro* proliferation of mesenchymal GICs was suppressed by a neutralizing antibody against APRIL in both U3031MG and U3054MG cells (Fig. 5H). The clinical significance of *APRIL* expression was analyzed using the TCGA dataset. The expression levels of *APRIL* were found to correlate with lack of *IDH* mutations, low G-CIMP expression, and poor prognosis of GBM patients (Fig. 5I-K).

TACI (also known as TNFRSF13B) and BCMA (TNFRSF17) are known as receptors for APRIL (*37*). However, analysis of the TCGA dataset revealed that the expression levels of *TACI/TNFRSF13B* or *BCMA/TNFRSF17* are very low in both normal and GBM tissues (fig. S4A). Moreover, flowcytometric analysis showed that expression levels of TACI and BCMA were not detectable or very low in the mesenchymal GICs, i.e., U3031MG and U3054MG cells, as well as in non-mesenchymal GICs, i.e., U3047MG and U3017MG cells used in the present study (fig. S4B, C). These findings strongly suggest that among the three TNF superfamily proteins which have been shown to interact with HVEM, APRIL binds to and activates HVEM in GBM cells in an autocrine fashion.

### HVEM inhibition sensitizes mesenchymal GICs to temozolomide and erlotinib

To investigate the effect of HVEM on the treatment with temozolomide or the epidermal growth factor receptor (EGFR) kinase inhibitor erlotinib, wild-type and HVEM-inactivated mesenchymal GICs (U3054MG-hCas9), as well as wild-type and HVEM-overexpressing non-mesenchymal GICs (U3017MG), were treated with temozolomide or erlotinib (Fig. 6A-D). HVEM inactivation sensitized mesenchymal GICs to temozolomide and erlotinib, while HVEM-expressing GICs exhibited enhanced resistance to these drugs. Moreover, both temozolomide and erlotinib efficiently induced apoptosis in HVEM-inactivated mesenchymal GICs (Fig. 6E), suggesting that blocking the HVEM function results in increased sensitivity of mesenchymal GICs to treatment with temozolomide or erlotinib.

**Fig. 6.**
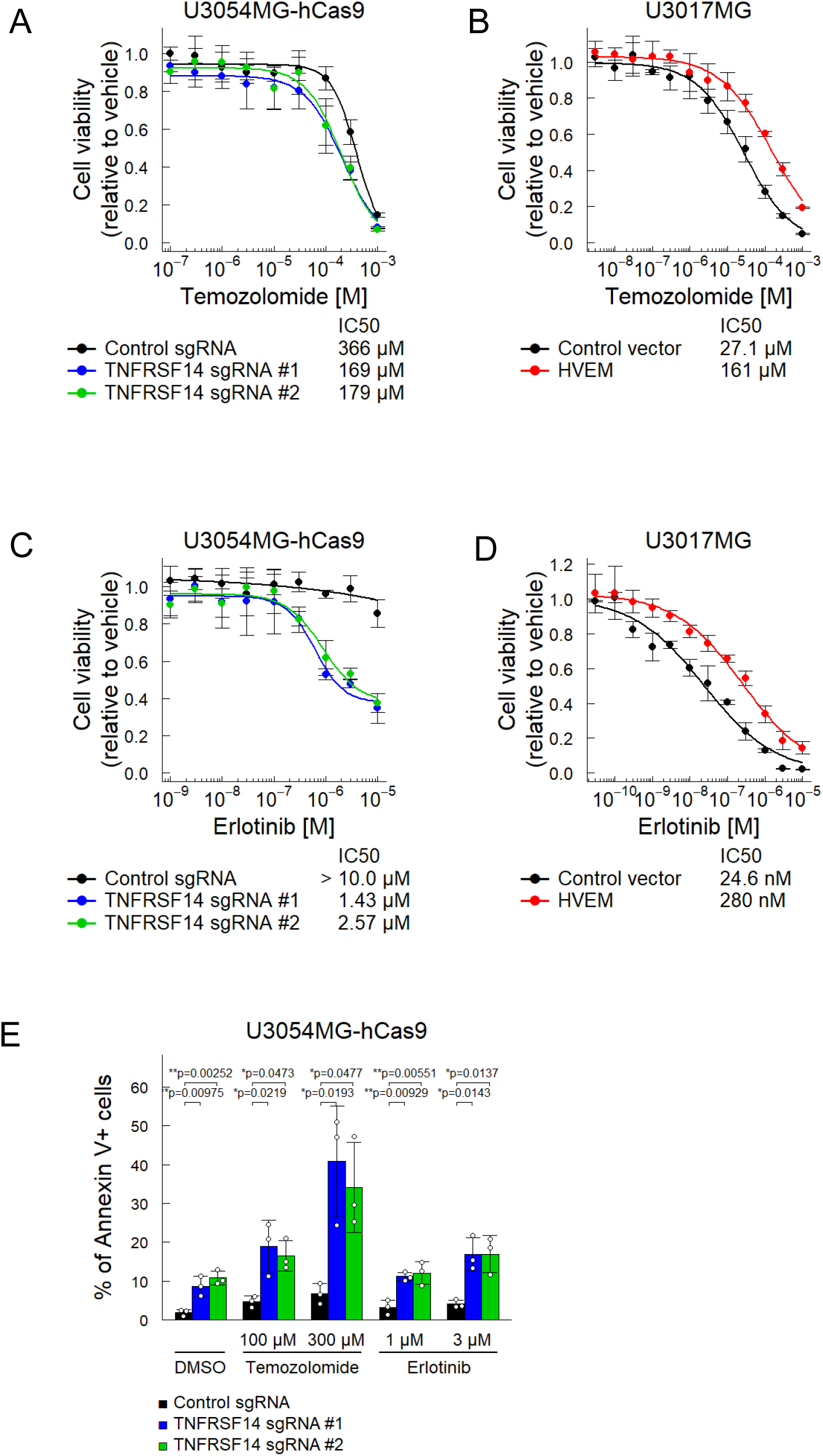
Effects of *HVEM/TNFRSF14* expression on sensitivity of GICs to anticancer drugs. (**A-D**) Effects of temozolomide (**A, B**) and erlotinib (**C, D**) on cell proliferation of *HVEM*-inactivated mesenchymal GICs (**A, C**) or *HVEM*-overexpressing non-mesenchymal GICs (**B, D**). GICs were treated for 7 days with temozolomide or erlotinib. (**E**) Apoptosis was monitored by annexin V staining. *HVEM*-inactivated mesenchymal GICs (U3054MG-hCas9) were treated for 7 days with indicated concentrations of temozolomide or erlotinib. Quantification data of annexin V-positive cells are shown as mean ± SD (n=3 biological replicates; \**P*<0.05, \*\**P*<0.01, \*\*\**P*<0.001; Tukey’s HSD test).

### An anti-human HVEM nanobody inhibits GBM invasion and proliferation

Camelids naturally produce heavy-chain-only antibodies. The variable domains of these antibodies known as nanobodies (also known as variable domains of heavy chain of heavy-chain-only antibodies or VHHs) are approximately 15 kDa in size, and have attracted considerable attention for clinical applications (*38*-*40*). To generate an anti-human HVEM nanobody, HVEM protein was immunized in alpacas. This approach yielded a nanobody that effectively inhibited the proliferation of U3054MG cells. An organotypic invasion assay revealed that the generated HVEM nanobody attenuated invasion of mesenchymal U03031MG and U3054MG cells (Fig. 7A). To investigate the effect of the anti-human HVEM nanobody on tumor progression *in vivo*, mesenchymal GICs were injected intracranially into nude mice. The anti-human HVEM nanobody was then administrated intraperitoneally every day after tumor engraftment. Inhibition of HVEM by the nanobody suppressed intracranial tumor proliferation (Fig. 7B), and prolonged the survival of mice bearing tumors (Fig. 7C).

**Fig. 7.**
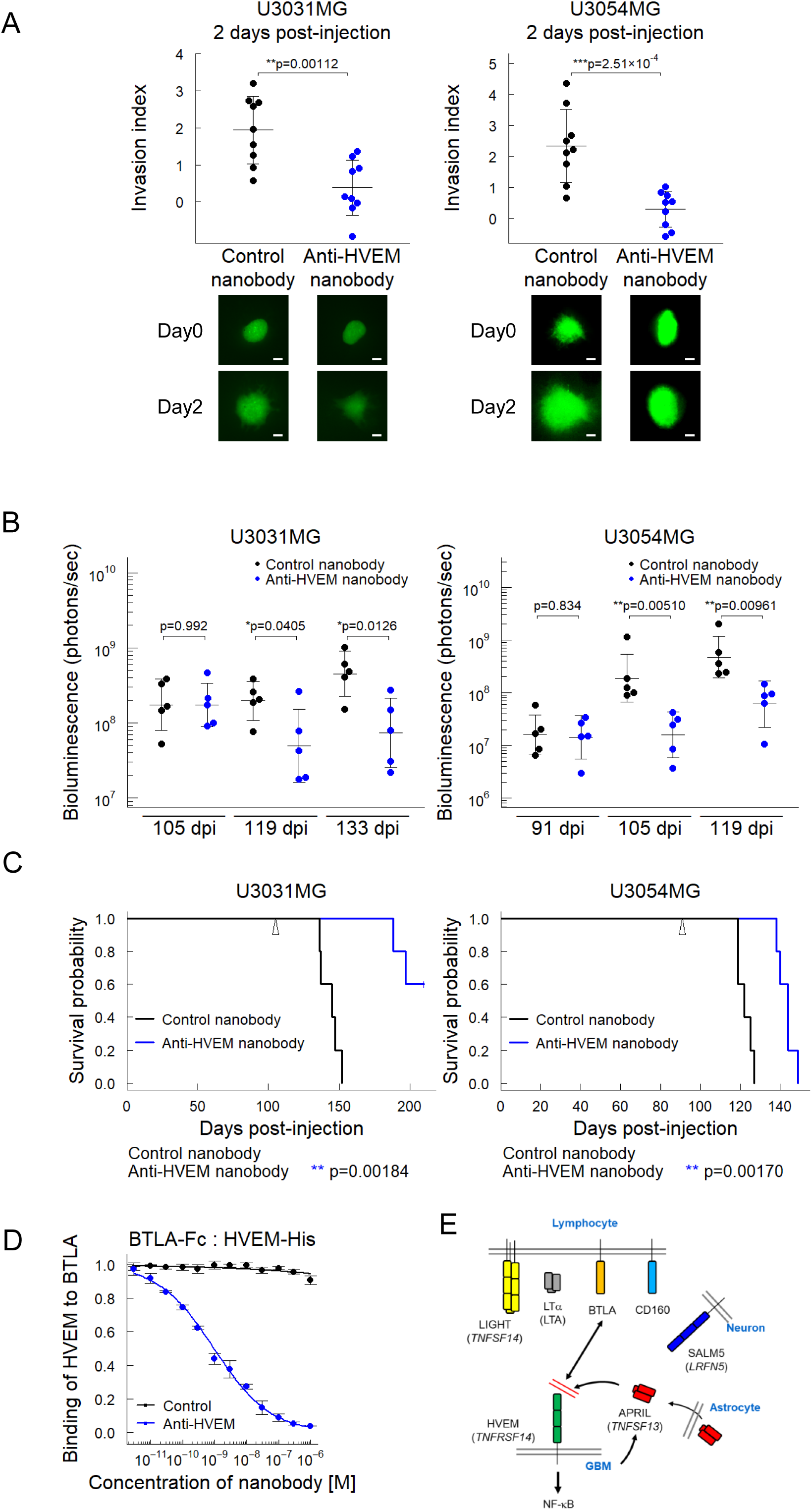
Effects of an anti-human HVEM nanobody on tumor progression of mesenchymal GICs. (**A**) Effects of the anti-human HVEM nanobody on invasion of mesenchymal GICs (U3031MG and U3054MG). Nanobodies were used at 1 μM. Fluorescence images with median invasion index in each group are demonstrated as representative images. Quantification data are shown as mean ± SD (n=9 biological replicates; \*\**P*<0.01, \*\*\**P*<0.001; two-tailed unpaired Student’s t-test). Scale bar: 200 μm. (**B, C**) *In vivo* effects of the anti-human HVEM nanobody on tumor progression of mesenchymal GICs. Mesenchymal GICs were injected intracranially into nude mice (n=5 mice per group). At 105 days (U3031MG) or 91 days (U3054MG) after tumor engraftment, the tumor size was quantified by bioluminescence imaging, and the mice were divided into two groups such that the amount of luminescence was equalized. Then, the nanobodies were administrated intraperitoneally at 10 mg/kg/day every day. Bioluminescence was quantified (\**P*<0.05, \*\**P*<0.01; two-tailed unpaired Student’s t-test) (**B**). Survival curves of mice bearing tumors (\*\**P*<0.01; two-tailed log-rank test) (**C**). (**D**) Inhibition of the interaction between HVEM and BTLA by the anti-human HVEM nanobody, as determined by an ELISA assay (n = 3, independent experiments). Binding of HVEM to BTLA was expressed as a ratio relative to the binding amount in the absence of the nanobody. (**E**) Dual inhibition of the anti-human HVEM nanobody, blocking the APRIL-HVEM axis as well as the HVEM-BTLA axis.

To exclude the possibility that the observed effects of the anti-human HVEM nanobody were caused by components present in the antibody preparations derived from mammalian cells, the gene encoding the anti-human HVEM nanobody or an anti-GFP nanobody was lentivirally introduced into U3054MG cells (fig. S5A). We confirmed that the anti-HVEM nanobody produced by the U3054MG cells efficiently bound to the HVEM protein ectopically expressed on human HEK293T cells (fig. S5B). The U3054MG cells expressing the anti-human HVEM nanobody exhibited significantly reduced proliferative capacity, as well as sphere-forming and invasive abilities, compared with control cells expressing the anti-GFP nanobody (fig. S5C-E).

### The anti-human HVEM nanobody prevents interaction between HVEM and BTLA

The extracellular domain of HVEM comprises of four cysteine-rich domains (CRDs), of which BTLA is known to bind to the most N-terminal domain, CRD1, while LIGHT binds to the CRD2 and 3 (*41*). Because of structural differences between the CRDs of human and mouse HVEM proteins, human HVEM does not interact with mouse-derived ligands, such as Btla, Cd160 and Salm5 (*42*, *43*). Based on this species specificity, we generated HEK293T cells expressing chimeric HVEM proteins containing either human- or mouse-derived CRD1, 2, and 3. We found that the anti-human HVEM nanobody recognized chimeric receptors containing human CRD1, but not those containing mouse CRD1 (fig. S6). Since BTLA binds to CRD1 of HVEM, we investigated whether the anti-human HVEM nanobody interferes with the interaction between BTLA and HVEM. As shown in figure 7D, the anti-human HVEM nanobody effectively blocked the interaction between BTLA and HVEM.

## DISCUSSION

### Multiple ligands interact with HVEM

We demonstrate that HVEM is highly expressed in GBM of the mesenchymal subtype. HVEM is a type I transmembrane protein expressed on certain tissues and cells, including T cells, B cells, natural killer cells, dendritic cells, hematopoietic cells, and non-hematopoietic cells (*26*, *27*, *44*). HVEM is well known as an immune checkpoint molecule with co-stimulatory and co-inhibitory functions; HVEM suppresses T cell immune response through its interaction with immunoglobulin-superfamily ligands, such as BTLA or CD160, whereas HVEM activates immune response upon binding to LIGHT. SALM5 is a synaptic adhesion molecule, which exhibits synaptogenic activities through binding to the leukocyte common antigen-related receptor protein tyrosine phosphatases (LAR-PTPs), including PTPδ (*45*-*47*). In addition, SALM5 acts as a ligand for HVEM to limit inflammatory responses in the central nervous system (CNS) (*43*). In cancer, elevated expression of *HVEM* is associated with poor prognosis in melanoma, esophageal, gastric, colon, breast, and non-small cell lung cancer (*48*-*52*).

The TNF superfamily ligand LIGHT induces trimerization of HVEM, leading to activation of NF-κB and JNK signaling pathways (*26, 53*, *54*). NF-κB signaling plays a crucial role in the regulation of cell proliferation and angiogenesis, and is critically involved in the development of cancer (*37*).

Among the genes encoding the known ligands for HVEM, increased expression of *LIGHT* was observed only in a subset of GBM tissues, whereas the expression of genes encoding other ligands, including *LTA,* were found to be very low in GBM tissues (Fig. 5B). *SALM5* was expressed in human normal brain tissues, but its expression was markedly reduced in human brain tumors. Furthermore, sandwich ELISA assays revealed that GBM cells do not produce detectable levels of either LIGHT or LTα (fig. S2B). Collectively, these findings suggest that ligands other than LIGHT or LTα are responsible for HVEM activation and the induction of the mesenchymal phenotype of GBM.

### APRIL is a ligand for HVEM in GBM

Deshayes and colleagues reported that APRIL stimulates the proliferation of certain human GBM cell lines, nerve progenitors, embryonic and adult astrocytes, cultured under serum-containing conditions (*55*). The GBM cells examined in their study included T98G, SF126, SNB19, U399MG, and U373MG cells. APRIL is known to bind to TACI and BCMA, which also serve as receptors for BAFF (TNFSF13B) (*56*). Although APRIL was able to bind to these cells, BAFF failed to bind to most of them. Based on these observations, the authors proposed the existence of APRIL-specific receptor(s) on the surface of GBM cell lines, nerve progenitors, and embryonic and adult astrocytes.

Here, we found that APRIL acts as a ligand for HVEM on GICs. Genetic disruption or antibody-induced inhibition of HVEM inhibited *in vivo* proliferation of human mesenchymal GICs in a mouse model. This finding suggests that tumor-intrinsic HVEM is activated by factors secreted by mesenchymal GICs; we demonstrated by a sandwich ELISA assay that APRIL is produced by mesenchymal as well as by non-mesenchymal GICs (Fig. 5C). APRIL is produced mainly by astrocytes in human brain, but by microglia in mouse brain (fig. S2A) (*57*). Moreover, the APRIL level was reported to be increased in serum of patients with brain tumors (*58*). Collectively, these findings suggest that APRIL binds to HVEM and acts on mesenchymal GBM cells to induce NF-κB signaling (Fig. 7E).

### Therapeutic potentials of nanobodies for treatment of GBM

Antibody-based therapies are widely used in the treatment of various cancers, because of their high specificity, low incidence of adverse effects, and long half-life in the circulation. However, conventional antibodies are relatively large (∼150 kDa), which limits their penetration into tumor tissues. Although camelid-derived nanobodies (∼15 kDa) have a short half-life in the circulation (less than 12 hours), they exhibit high tissue penetration, including penetration into the brain (*38*-*40*). To overcome their short half-life, strategies such as attachment to IgG Fc region or serum albumin-binding proteins have been employed. In this study, we generated a camelid-derived nanobody to human HVEM, and found that it suppressed the invasion of GICs in organotypic cultures and inhibited tumor formation in an orthotopic xenograft model (Fig. 7A-C). Because the amino acid sequences of HVEM are highly divergent between human and mouse, it is critical to generate antibodies specific to human HVEM for clinical application.

The blood-brain barrier (BBB) restricts the passage of high-molecular weight therapeutics, including conventional antibodies, into the brain, and extensive efforts have been made to overcome this limitation. In GBM tissues, structural alterations of the BBB are frequently observed, leading to the formation of a more permeable interface, referred to as blood-tumor barrier (BTB). Although certain drugs can penetrate GBM tissues through the BTB and the integrity of the BTB is highly heterogeneous across different regions of the tumor (*59*), development of strategies that further enhance drug penetration is critical for the effective treatment of GBM (*60*). In the present study, due to its short half-life in circulation, the nanobody was administered via daily intraperitoneal injections. For clinical use, it will thus be essential to prolong the half-life of the nanobody while preserving its ability to penetrate brain tissues.

### Possible effects of HVEM inhibition on immune regulation in GBM

In addition to members of the TNF superfamily, HVEM also binds to immunoglobulin-superfamily ligands, *i.e.* BTLA and CD160. The TNF superfamily member LIGHT induces trimerization of HVEM, whereas BTLA and CD160 interact with HVEM in monomeric forms (*41*, *61*). We observed that the anti-human HVEM nanobody efficiently inhibited the interaction between HVEM and BTLA via binding to CRD1 of HVEM (Fig. 7D). BTLA is an inhibitory factor in immune checkpoints and is known to negatively regulate T-cell-mediated immune responses (*62*, *63*). Moreover, targeting the BTLA-HVEM axis has been shown to enhance antitumor immunity, and promising results have been reported in anticancer therapy targeting this pathway (*44, 64*, *65*). Recent studies have demonstrated that immune checkpoint blockade through inhibition of the HVEM-BTLA interaction enhanced immune response upon combination therapy with anti-PD-1 antibody (*66*, *67*). In accordance, we found that expression levels of *CD274* and *PDCD1LG2,* encoding PD-L1 and PD-L2, respectively, are elevated in mesenchymal GBMs (Fig. 1A). Thus far, immune checkpoint inhibitors, including anti-PD-1 or anti-PD-L1 antibodies, have not demonstrated substantial clinical effects in patients with GBM (*68*). These findings suggest that combination strategies simultaneously targeting PD-1 and HVEM may enhance therapeutic outcomes in GBM.

HVEM is highly expressed in GBM and is associated with poor patient prognosis (*69*). Recently, Han and colleagues demonstrated that HVEM/TNFRSF14 expression is upregulated by interferon-γ signaling. Consistent with our findings, they showed that disruption of HVEM expression reduced the tumorigenicity of GBM cells, although the specific ligand(s) for HVEM in GBM tissues were not identified. They further demonstrated that HVEM induces the activation of NF-κB signaling, and that administration of anti-mouse HVEM antibody enhances the therapeutic efficacy of anti-PD-L1 in mouse orthotopic glioma model (*70*).

### Limitations of the present study

In this study, immune-compromised mice were used to evaluate the *in vivo* effects of HVEM on GBM cells. The tumor microenvironment plays a critical role in GBM progression, particularly in mesenchymal GBM, where tumor-associated macrophages substantially influence tumor behavior. In addition, the amino acid sequences of HVEM and its ligands differ markedly between species. Therefore, the effects of HVEM observed in the present study likely reflect tumor-intrinsic mechanisms, and the contributions of cells within the tumor microenvironment were not fully evaluated. Nevertheless, analysis of the TCGA dataset demonstrated that both APRIL and HVEM are highly expressed in tumor tissues compared to other ligands and receptors (Fig. 5B and fig. S4A). Further studies are required to elucidate the role of the tumor microenvironment in modulating HVEM signaling in GBM.

The nanobody described in the present study appears to exert dual effects on mesenchymal GBM cells; it inhibits tumor cell growth and invasion of GBM cells driven by the APRIL-HVEM-NF-κB axis, and alleviates immune suppression by blocking the HVEM-BTLA axis on immune cells (Fig. 7E). Tumor intrinsic HVEM is thus a clinically interesting target for treatment of mesenchymal GBM not only as immune checkpoint blockade therapy. Collectively, these findings indicate that blockade of the function of HVEM can provide clinical benefits with synergistic effects in combination with immune checkpoint inhibitor(s) to inhibit GBM progression.

## MATERIALS AND METHODS

### Study design

This study was designed to identify therapeutic target(s) in mesenchymal GBM and to develop effective antibody therapeutics against those molecule(s). To this end, candidate molecules were identified by searching the HGCC Resource of human GBM cells cultured under serum-free conditions (*24, 25*; https://www.hgcc.se/), and their expression profiles and associations with prognosis in GBM patients were analyzed using the TCGA dataset (Fig. 1). Focusing on HVEM as a therapeutic candidate, *HVEM* gene down-regulation by knockdown or knockout, or artificial overexpression in GBM cells was performed. After confirming HVEM protein expression by flowcytometry analysis, the role of HVEM in GBM cells was analyzed by cell proliferation, sphere formation, and organotypic cell invasion assays. Furthermore, orthotopic transplantation experiments using nude mice were conducted to evaluate tumor growth in mice and overall survival (Figs. 2–4). BMP-4 was examined as a regulator of HVEM expression (fig. S1).

The expression of HVEM binding partners was investigated using TCGA and other datasets, and their production in cultured GBM cells was assessed by sandwich ELISA assays. APRIL was then evaluated for its binding ability to HVEM using pulldown and NF-κB reporter assays. The roles of APRIL in GBM cells were examined by cell proliferation, sphere formation, and organotypic invasion assays upon knockdown of the *APRIL* gene. In addition, the expression profile of APRIL and its association with prognosis in GBM patients were validated using the TCGA dataset. The expression of known APRIL receptors in brain tumor cells was analyzed using TCGA dataset, and their expression in GBM cells was examined by flow cytometry (Fig. 5, figs. S2–S4). The role of HVEM in drug resistance of mesenchymal GBM cells was evaluated by knocking out or overexpressing HVEM in GBM cells, followed by analyss using cell viability and apoptosis induction assays (Fig. 6).

Considering HVEM as a promising therapeutic target for mesenchymal GBM, nanobodies against human HVEM were generated. Their effects on GBM cells were evaluated in terms of invasive capacity, as well as tumor growth in the mouse brain and overall survival. In addition, GBM cells were transduced with the nanobody gene to confirm the nanobody protein expression, followed by analyses using cell proliferation, sphere formation, and organotypic invasion assays. Furthermore, the binding epitope of the nanoantibody on HVEM was identified, and an ELISA assay was used to determine whether the nanobody also affected the binding between BTLA and HVEM (Fig. 7, figs. S5–S6).

Detailed experimental procedures are described below and in the Supplementary Materials and Methods. The experiments were conducted at the University of Tokyo and Uppsala University in accordance with the experimental guidelines established by each institution.

### Cell culture and reagents

Human GBM cells, classified into the four GBM subtypes according to the data (GSE72217), were obtained from the HGCC Resource (*24*, *25*). All the GBM cells used in this study were obtained in accordance with informed patient consent and with approval from the Uppsala ethical review board (2007/353). GBM cells were cultured in DMEM/F12 (Thermo Fisher Scientific) and Neurobasal medium (Thermo Fisher Scientific), supplemented with B-27 supplement (Thermo Fisher Scientific), N-2 supplement (Thermo Fisher Scientific), 20 ng/mL of epidermal growth factor (EGF, PeproTech) and 20 ng/mL of basic fibroblast growth factor (bFGF, PeproTech). H9 human embryonic stem cell (hESC)-derived human neural stem cells (Thermo Fisher Scientific) were cultured in KnockOut DMEM/F12 (Thermo Fisher Scientific), supplemented with StemPro neural supplement (Thermo Fisher Scientific), EGF (20 ng/mL) and bFGF (20 ng/mL).

### Production and purification of anti-human HVEM nanobodies

For the generation of nanobodies against human HVEM, immunization of alpacas and collection of the blood were conducted at ARK Resource, Co. (Kumamoto, Japan), and genes encoding anti-human HVEM nanobodies were isolated. DNA sequences encoding anti-GFP or anti-HVEM nanobodies tagged with 6xHis at the C-terminus were cloned into pcDNA3.4-TOPO (Thermo Fisher Scientific). Using Expi293 Expression System (Thermo Fisher Scientific), pcDNA3.4 plasmids were transfected into Expi293F cells. 6xHis-tagged nanobodies produced in cell cultures were purified with Ni-NTA resin, and subjected to endotoxin removal with ToxinEraser Endotoxin Removal Kit (#L00338, GenScript). Endotoxin levels in purified nanobodies were determined with ToxinSensor Chromogenic LAL Endotoxin Assay Kit (#L00350, GenScript).

### Intracranial proliferation assay

All animal experiments were approved by and carried out according to the guidelines of the Animal Ethics Committee of the Graduate School of Medicine, the University of Tokyo. A total of 1×10^5^ viable cells were injected over bregma 2 mm to the right of the sagittal suture and 3 mm below the surface of the skull of 5-week-old female BALB/c *nu/nu* mice (SLC). *In vivo* bioluminescent imaging was conducted on NightOWL LB981 NC-100T (Berthold technologies) after *firefly* luciferin (Promega) was injected intraperitoneally into mice. Following the rules of the Animal Committee of the University of Tokyo, mice were monitored until they exhibited reduction of body weight by more than 20% or neurological signs, such as hemiparesis and domehead.

### Statistical analysis

Statistical analyses (Student’s t-test, Welch’s t-test, Mann-Whitney U test, two-way ANOVA and log-rank test) were performed by statistical software R (http://www.R-project.org). The Cancer Genome Atlas (TCGA) was downloaded from the TCGA data portal (https://tcga-data.nci.nih.gov) and the NIH’s Genomic Data Commons (https://api.gdc.cancer.gov). Expression data (GSE72217, GSE52564 and GSE73721) was downloaded from NCBI Gene Expression Omnibus (https://www.ncbi.nlm.nih.gov/geo/) and HGCC (https://www.hgcc.se/).

## Supplementary Materials

### The PDF file includes

Matrials and Methods

Figs. S1 to S6

Tables S1 to S3

## Acknowledgments

We express our sincere appreciation to the patients and their families for their support and for donating materials to the HGCC biobank. We also acknowledge the HGCC team for their invaluable contributions in collecting and providing the patient-derived GIC cultures utilized in this study. We thank Drs. Takehisa Matsumoto, Mikako Shirouzu (RIKEN), Koji Tamada (Yamanashi University), Yoshitaka Narita (National Cancer Center, Japan), Shota Tanaka (Okayama University), Tomoko Ozawa (UCSF), and Masato Morikawa (Teikyo University) for valuable suggestions and discussion.

## Funding

Swedish Cancer Society grant 100452 (KM)

Swedish Cancer Society grant 222363 (C-HH)

Swedish Cancer Society grant 243486 (BW)

Swedish Research Council grant 2024-03002 (C-HH)

The Japan Society for the Promotion of Science (JSPS) KAKENHI Innovative Area on Integrated Analysis and Regulation of Cellular Diversity grant JP17H06326 (KM)

JSPS KAKENHI Scientific Research (S) grant JP15H05774 (KM)

JSPS KAKENHI Scientific Research (S) grant JP23H05486 (KM)

JSPS TR-SPRINT grant JP19lm0203003 (KM)

Japan Agency for Medical Research and Development (AMED)

Practical Research for Innovative Cancer Control grant JP21ck0106705 (KM)

AMED Practical Research for Innovative Cancer Control grant JP24ck0106914 (KM)

The University of Tokyo GAP Fund Program (KM)

## Author contributions

Conceptualization: RT, BW, CHH, KM

Methodology: RT, BW, CHH, KM

Investigation: RT

Visualization: RT

Funding acquisition: BW, CHH, KM

Project administration: RT, KM

Supervision: BW, CHH, KM

Writing – original draft: RT

Writing – review & editing: RT, BW, CHH, KM

## Data and materials availability

All data associated with this study are present in the paper or the Supplementary Materials and Methods. Amino acid sequence of the anti-human HVEM nanobody is shown as VHH#3 in WO 2020/138503. Further information and materials are available upon reasonable request under materials transfer agreements (MTAs).

