## Supplementary materials and figures for "Inhibition of HVEM suppresses growth and invasion of mesenchymal glioblastoma"

### SUPPLEMENTARY INFORMATION

#### SUPPLEMENTARY MATERIALS AND METHODS

##### Reagents

The antibodies used were as follows; PE-conjugated anti-TNFRSF14 antibody (#130-101-593, Miltenyi Biotec), PE/Cyanine7-conjugated anti-TACI/TNFRSF13B antibody (#311907, BioLegend), PE-conjugated anti-BCMA/TNFRSF17 antibody (#357503, BioLegend), PE-conjugated REA Control human IgG1 (#130-104-612, Miltenyi Biotec), PE/Cyanine7-conjugated control rat IgG2a (#400522, BioLegend), PE-conjugated control mouse IgG2a (#400212, BioLegend), anti-TNFSF13/APRIL (#MAB5860, R&D SYSTEMS), Isotype control mouse IgG1 (#MAB002, R&D SYSTEMS), HRP-conjugated anti-His tag antibody (#D291-7, MBL), HRP-conjugated anti-HA tag antibody (#M180-7, MBL), HRP-conjugated anti-human IgG Fc antibody (#ab98624, abcam), and Alexa Fluor 488-conjugated anti-His tag antibody (#291-A48, MBL). Temozolomide and erlotinib were purchased from MedChemExpress. BMP-4 was purchased from R&D SYSTEMS.

##### Lentiviral transduction

Genes or promoters were cloned into pENTR201 or pENTR4 vectors, and then recombination between pENTR and lentiviral vectors (CS-RfA-CMV-PuroR, CS-RfA-EF-PuroR, CSII-CMV-RfA, CS-RfA-EG or CS-RfA-CMV-hRluc) was catalyzed by LR clonase (Thermo Fisher Scientific), as previously described (*34*). DNA sequences encoding sgRNA were inserted into lentiCRISPR v2 vector. Lentiviral vector plasmids used are listed in table S1. Detailed sequences of short hairpin RNAs (shRNAs) and single guide RNAs (sgRNAs) are in table S2. The indicated lentiviral vector plasmids and lentivirus packaging vector plasmids (pCAG-HIVgp and pCMV-VSV-G-Rev) were transduced into HEK293FT cells (Thermo Fisher Scientific) with Lipofectamine 2000 (Thermo Fisher Scientific). Lentiviral particles were concentrated with Lenti-X Concentrator (Takara Bio) and then reconstituted with serum-free buffer.

##### Quantitative real-time PCR analysis

Total RNA was extracted from cells with RNeasy Mini Kit (Qiagen). Complementary DNA was synthesized with PrimeScript II First Strand cDNA Synthesis Kit (Takara Bio). Gene expression levels were quantified using Step One Plus Real-Time PCR Systems (Applied Biosystems) with FastStart Universal SYBR Green Master Mix (Roche), using the primer sets listed in table S3.

##### Sandwich ELISA

Cells were cultured for 24 hours, and then the cell supernatants were collected and filtrated using 0.45-μm filters. Expression levels of human APRIL and other potential HVEM ligands in the cell supernatants were quantified by immunoassays, i.e. Ancillary Reagent Kit 2 (#DY008, R&D SYSTEMS), Human APRIL/TNFSF13 DuoSet ELISA (#DY884B, R&D SYSTEMS), Human LIGHT/TNFSF14 Quantikine ELISA kit (#DLIT00, R&D SYSTEMS), and Human Lymphotoxin-alpha/TNF-beta DuoSet ELISA (#DY211, R&D SYSTEMS). Absorbance at 450 nm and 570 nm was measured on Model680 microplate reader (Bio-Rad) or Enspire (Perkin Elmer). To investigate effects of nanobodies on HVEM-BTLA interaction, recombinant human HVEM tagged with poly-histidine (#10334-H08H, Sino Biological) was immobilized on copper-coated immunoassay plates (#15146, Thermo Fisher Scientific). The immobilized HVEM was incubated with nanobodies, followed by incubation with human BTLA-Fc (#31043, PeproTech) for 1 hour at room temperature; the recombinant human BTLA-Fc that bound to the recombinant human HVEM was measured using HRP-conjugated goat anti-human IgG Fc.

**Flowcytometric analysis**
Cells were dissociated with StemPro Accutase Cell Dissociation Reagent (#A1110501, Thermo Fisher Scientific), and then stained with PE-conjugated anti-TNFRSF14, PE/Cyanine7-conjugated anti-TACI, PE-conjugated anti-BCMA, or isotype control antibodies. To detect reactivity of an anti-human HVEM nanobody tagged with 6xHis, cells were incubated with an anti-human HVEM nanobody and then stained with Alexa Fluor 488-conjugated anti-His antibody. Protein expression on the cell surface was analyzed using SH800S Cell Sorter (Sony) and the FlowJo software (Beckton Dickinson).

**Pulldown assay**
Dynabeads His-Tag Isolation & Pulldown (#10104D, Thermo Fisher Scientific) to which His-tagged human HVEM protein (#H10334-H08H, Sino Biological) or His tag peptide (#3310-205, MBL) were coupled, were mixed with HA-tagged human APRIL protein (#5860-AP/CF, R&D SYSTEMS), and His-tagged proteins were eluted following the manufacturer's instructions, denatured at 98°C for 5 min, and then subjected to sodium dodecyl sulfate-polyacrylamide gel electrophoresis (SDS-PAGE), followed by immunoblot analysis.

***In vitro* proliferation assay**
Cells were seeded on 96-well microplates and cultured for up to 7 days. Cell counting kit-8 (Nacalai Tesque) was used to determine the cell number, following the manufacturer's instructions. Absorbance at 450 nm and 595 nm was measured on Model680 microplate reader (Bio-Rad) or Enspire (Perkin Elmer).

**EdU incorporation assay**
Cells were cultured for 48 hours and then labeled for 2 hours with 10 μM of EdU. The cells were fixed by incubation in 3.7% formaldehyde for 15 minutes and then permeabilized by incubation in 0.5% Triton X-100 for 20 minutes. The incorporated EdU was detected by using EdU-Click 647 kit (#BCK-EDU647, Merck). Nuclei were stained with VECTASHIELD Antifade Mounting Medium with DAPI (#H-1200, Vector Laboratories). Cell images were captured using a BZ-X710 microscope (Keyence).

**Limiting dilution and sphere formation assay**
Cells were seeded in ultra-low attachment microplates (Corning) at a density of 1-100 cells/well and cultured for 7 days. Cell masses of a diameter more than 20 μm were considered spheres. Wells with or without spheres were counted in each condition.

**Organotypic invasion assay**
Murine brains were sliced at a thickness of 300-500 μm in coronal sections. The brain slices were cultured in 0.4-μm membrane inserts placed in 6-well plates. Human GBM cells were labelled by carboxyfluorescein succinimidyl ester (CFSE), and then cultured for 24 hours in ultra-low attachment microplates at a density of $1\times10^4$ cells/well to generate glioma spheres. CFSE-labeled spheres were placed on murine brain slices and cultured for 2 days. Fluorescence images were

obtained at day 0 and 2 using a BZ-X710 microscope, and fluorescent areas were then evaluated by ImageJ (https://imagej.net/ij/). Invasion areas were determined by subtraction of the fluorescent areas at day 0 from those at day 2.

**Bioluminescence-based reporter assay**

To measure NF-κB activity, *firefly* luciferase gene driven by minimal promoter (minP) with NF-κB responsive elements was transduced lentivirally into cells together with *renilla* luciferase gene driven by CMV promoter. Using Dual-Luciferase Reporter Assay System (E1980, Promega), activities of *firefly* and *renilla* luciferases were measured by Mithras LB940 (Berthold technologies) or Infinite 200 PRO plate readers (Tecan). *Firefly* luciferase activity was standardized by *renilla* luciferase activity to determine NF-κB activity.

**Apoptosis detection assay**

Cells were dissociated with StemPro Accutase Cell Dissociation Reagent (#A1110501, Thermo Fisher Scientific), and then stained with FITC Annexin V Apoptosis Detection Kit with 7-AAD (#640922, BioLegend). Apoptotic cells were analyzed by SH800S Cell Sorter (Sony) and the FlowJo software (Beckton Dickinson).

**Ligand stimulation of HVEM and determination of NF-κB activation**

For preparation of cells expressing HVEM and NF-κB reporter genes, CS-NF-κB-RE-minP-Luc2-CMV-hRluc in which the firefly luciferase gene (Luc2, Promega Corporation) was cloned downstream of the minimal promoter with the NF-κB responsive element and the Renilla luciferase gene (hRluc, Promega Corporation) cloned downstream of the CMV promoter was prepared. After the NF-κB responsive Luc2 gene and the constantly-expressed hRluc gene were introduced lentivirally into HEK293T cells, the cells were infected with a human HVEM expression lentiviral vector or an empty vector (target cells). Then, LIGHT, APRIL and SALM5 were each expressed lentivirally in HEK293T cells without the NF-κB reporter gene to prepare cells expressing soluble or membrane-bound forms of ligands (effector cells). As control cells, cells expressing the NF-κB reporter gene, but neither expressing HVEM nor the ligands, were prepared (Empty). The HVEM-expressing cells and the ligand-expressing cells were co-cultured, and the activity of NF-κB in the HVEM-expressing cells was evaluated using Dual Luciferase Reporter Assay Kit (Promega).

#### SUPPLEMENTARY FIGURES

**Fig. S1. Effects of BMP-4 on GICs of mesenchymal subtype.** (**A, B**) Growth curves (**A**) and sphere formation (**B**) of mesenchymal GICs (U3024MG, U3031MG, and U3054MG) in the presence or absence of 30 ng/mL BMP-4. Data are shown as mean ± SD (n=3 biological replicates; ***P*<0.01, ****P*<0.001; two-tailed unpaired Student's t-test for cell proliferation assay, two-way ANOVA for sphere formation assay). (**C**) Quantitative RT-PCR analysis of *HVEM/TNFRSF14* in GICs treated or not with BMP-4 for 48 hours. Expression levels of *HVEM* in the mesenchymal GICs (U3024MG and U3031MG) which responded to BMP-4 stimulation (**A, B**), were analyzed.

**Fig. S2. Analysis of LTα and LIGHT concentrations in mesenchymal and non-mesenchymal GICs.** (**A**) A heatmap of the differentially expressed genes encoding APRIL and known ligands for HVEM in the human (*left*) and mouse (*right*) brain. Expression data from GSE73721 and GSE52564 were re-analyzed, respectively. (**B**) Expression levels of LTα and LIGHT in hNSCs and GICs from the HGCC Resource were determined by sandwich ELISA. Expression levels of LTα and LIGHT were below the detection limit (not detected; n.d.).

**Fig. S3. NF-κB activation by HVEM ligands.** (**A**) HEK293T cells expressing HVEM (target cells) were co-cultured with HEK293T cells expressing a membrane-bound or soluble form of ligand (APRIL, LIGHT or SALM5) (effector cells). Note that APRIL is produced exclusively as a soluble form (*36*) (**B, C**) Induction of NF-κB activity in the target cells co-cultured with the effector cells expressing various membrane-bound (**B**) and soluble forms of ligands (**C**). Data is shown as mean ± SD for NF-κB relative activity in HEK293T cells expressing HVEM or control vectors. E:T ratio indicates the ratio of effector cells and target cells.

**Fig. S4. Expression of receptors for APRIL in the mesenchymal GBM.** (**A**) Expression levels of mRNAs encoding known receptors for APRIL (*BCMA/TNFRSF17* and *TACI/TNFRSF13B*) and *HVEM/TNFRSF14* in normal brain tissue and GBM in the TCGA dataset. Expression level of mRNA for HVEM in normal brain tissues shown in Fig. 1C is shown here for comparison (****P*<0.001, **P*<0.05, not significant (*P*>0.05); two-tailed Kruskal-Wallis test). (**B, C**) Cell surface expression of HVEM, TACI, and BCMA in the mesenchymal GICs (**B**) and that of TACI and BCMA in the non-mesenchymal GICs (**C**) were evaluated by flowcytometric analysis.

**Fig. S5. Effects of gene introduction of the anti-human HVEM nanobody in U3054MG cells.** (**A**) Immunoblot analysis of the expression of His-tagged nanobodies after introducing the gene of alpaca anti-human HVEM nanobody or an anti-GFP nanobody (negative control) into U3054MG cells. (**B**) Binding of anti-HVEM nanobody or anti-GFP nanobody produced by the U3054MG cells to HVEM. Culture medium of U3054MG cells obtained in (**A**) was added to HEK293T cells expressing HVEM or not (control), and the binding of the nanobodies to HVEM was analyzed using flowcytometry. (**C, D**) Growth curves (**C**) and sphere forming ability, as determined by limiting dilution assay (**D**), of U3054MG cells expressing the anti-human HVEM nanobody or the anti-GFP nanobody. Data are shown as mean ± SD (n=3 biological replicates; ***P*<0.01, ****P*<0.001; two-tailed unpaired Student's t-test) (**C**) and as mean ± SD (n=3 independent experiments; ****P*<0.001; two-way ANOVA) (**D**). (**E**) Invasiveness of U3054MG cells expressing the anti-human HVEM nanobody or the anti-GFP nanobody was evaluated by organotypic invasion assay. Fluorescence images with median invasion index in each group are demonstrated as representative images (n=9 biological replicates; ***$P$<0.001; two-tailed unpaired Student's t-test). Scale bar: 200 μm.

**Fig. S6. Binding of the anti-human HVEM nanobody to chimeric HVEM proteins.** Three CRDs of human HVEM were replaced with the corresponding regions of mouse CRDs. Binding of the anti-human HVEM nanobody to HEK293T cells expressing chimeric HVEM proteins was analyzed using flowcytometry.

Supplementary Figure S1.

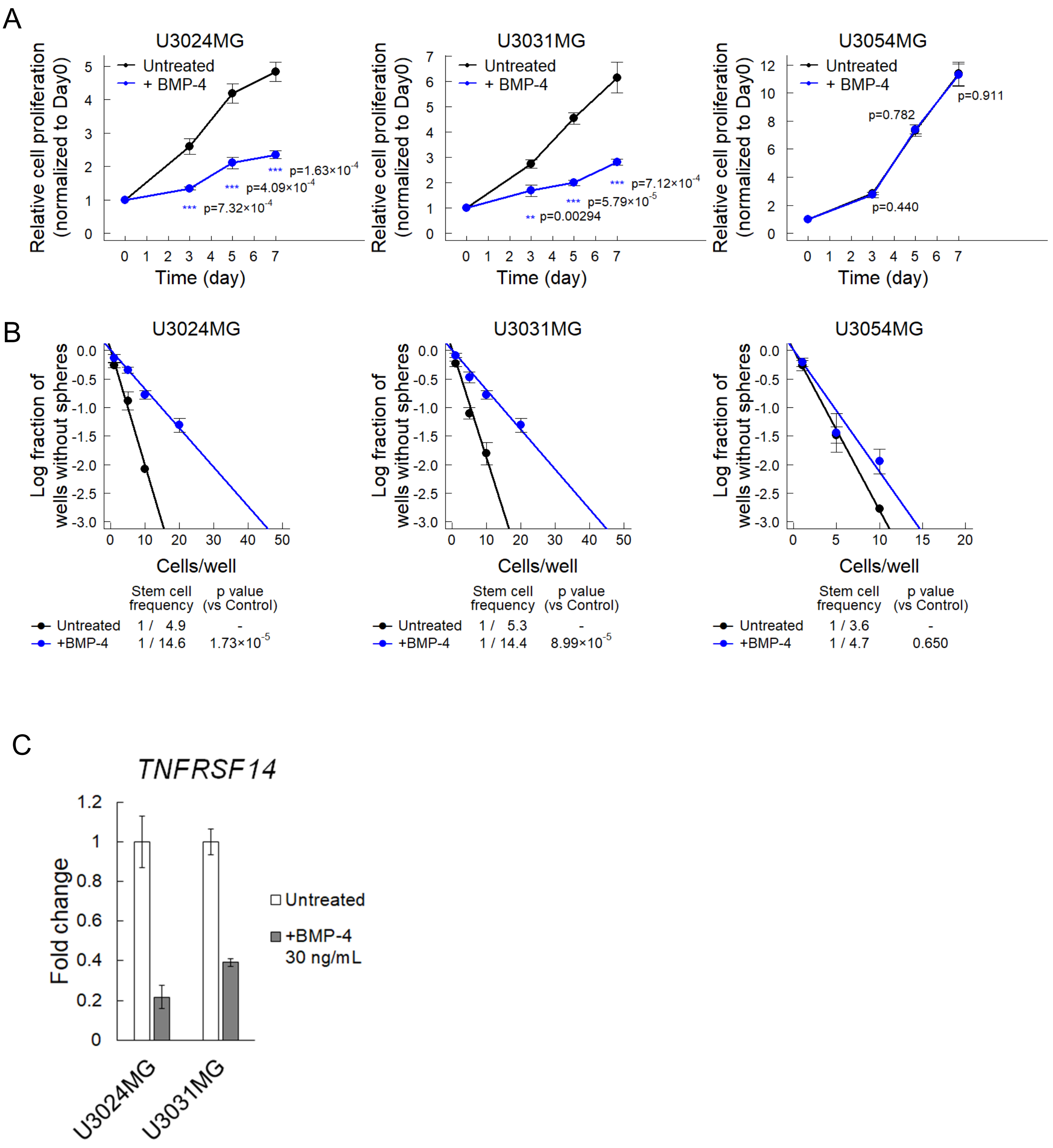

Supplementary Figure S2.

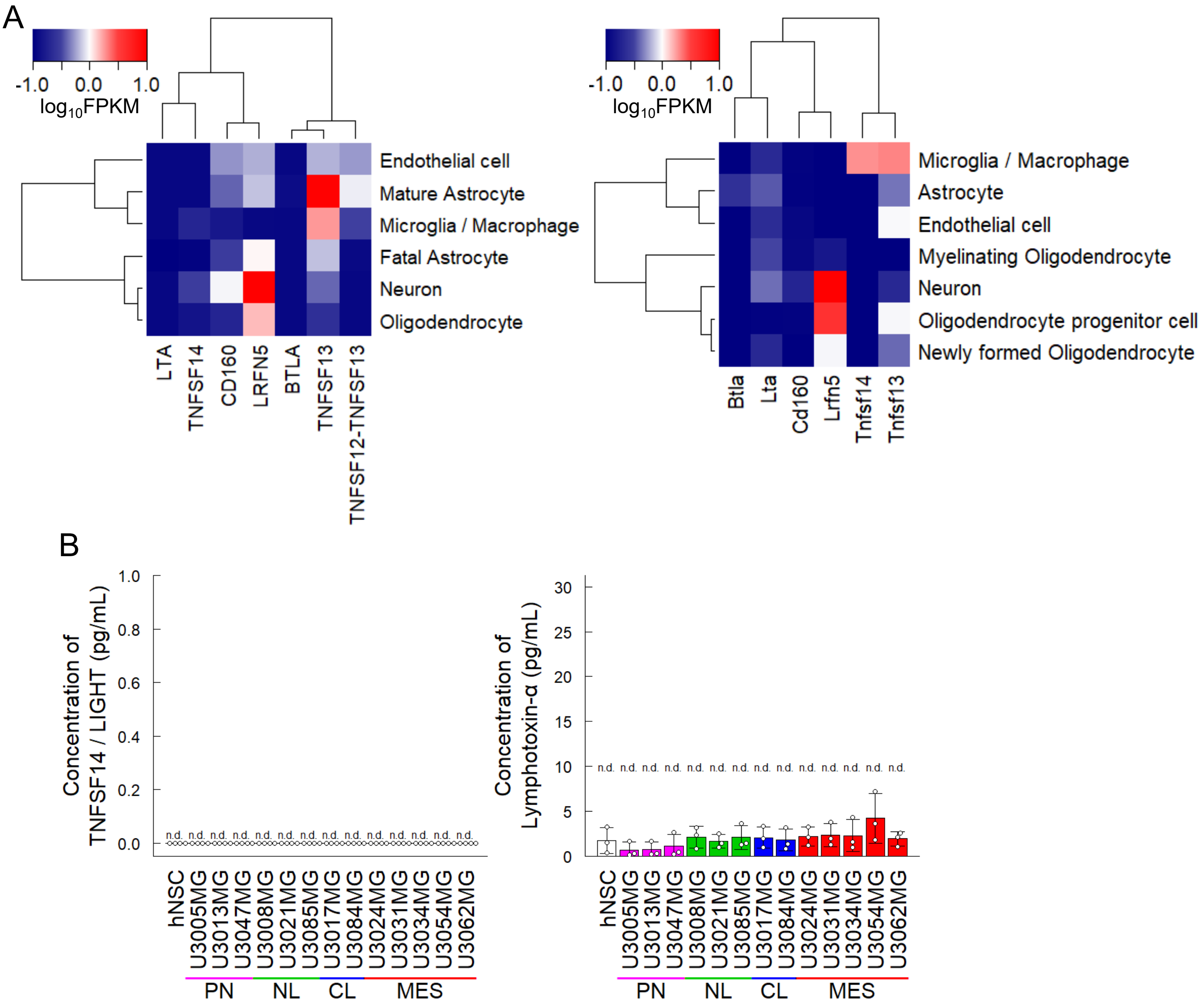

Supplementary Figure S3.

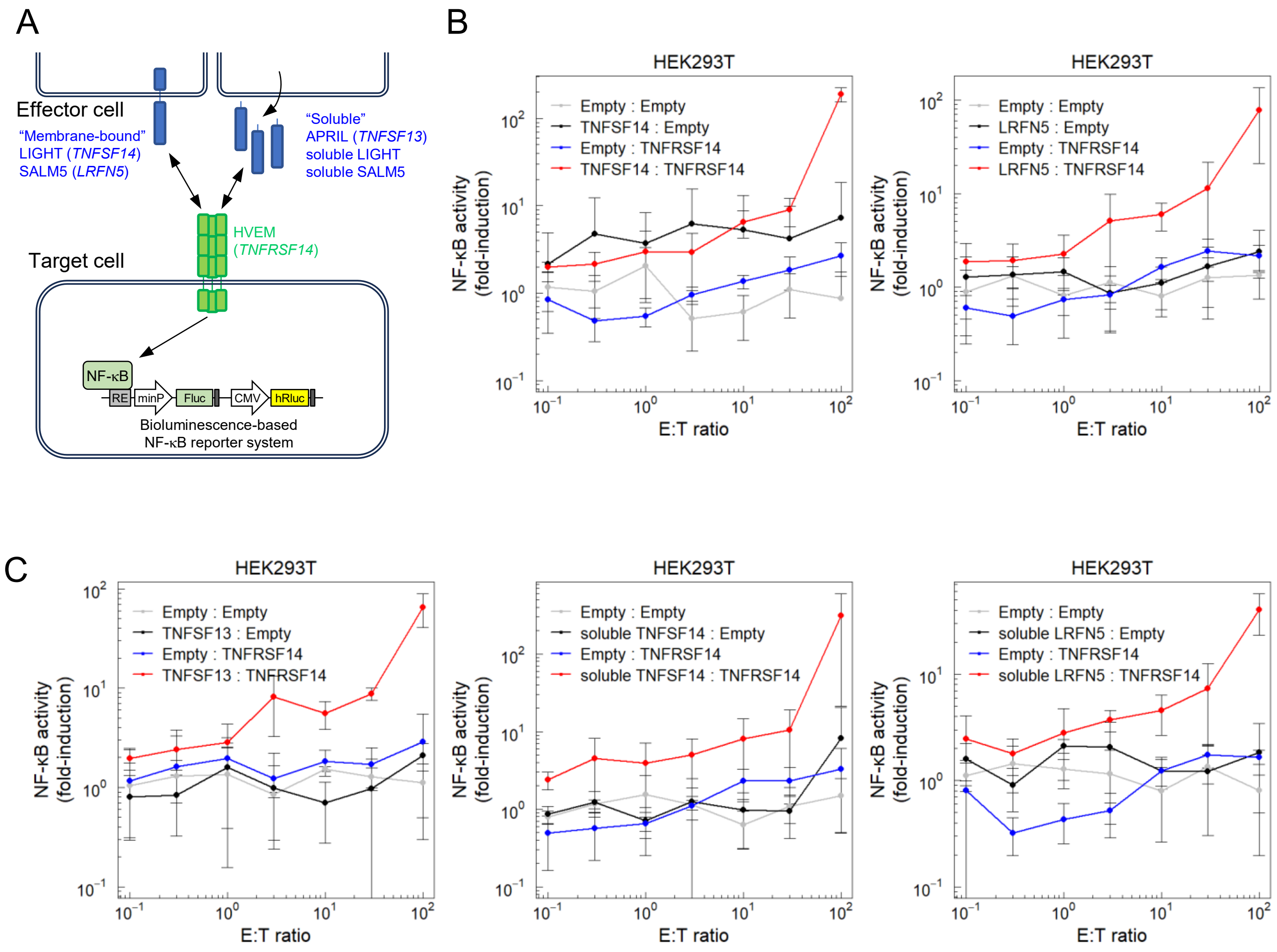

Supplementary Figure S4.

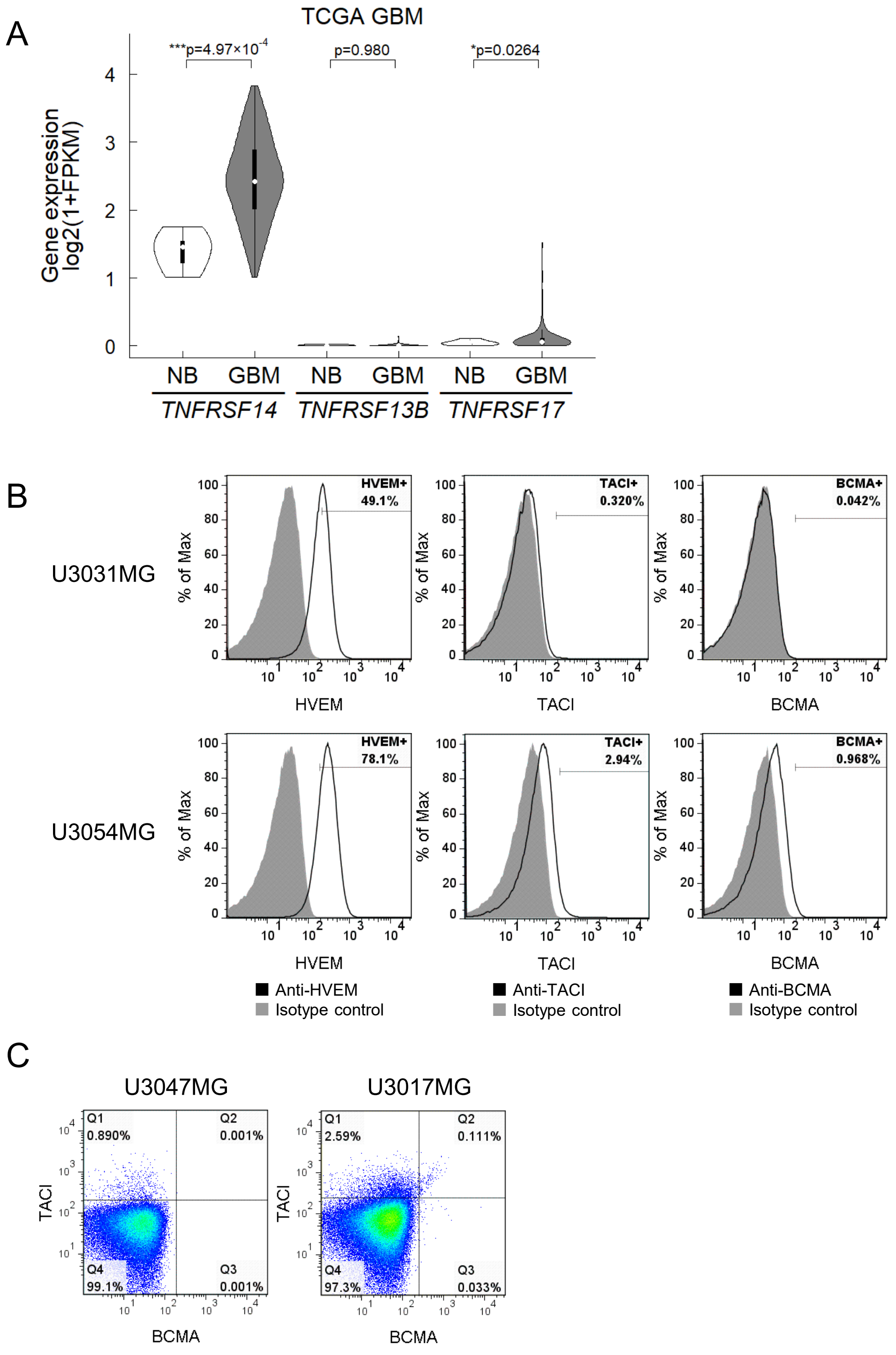

Supplementary Figure S5.

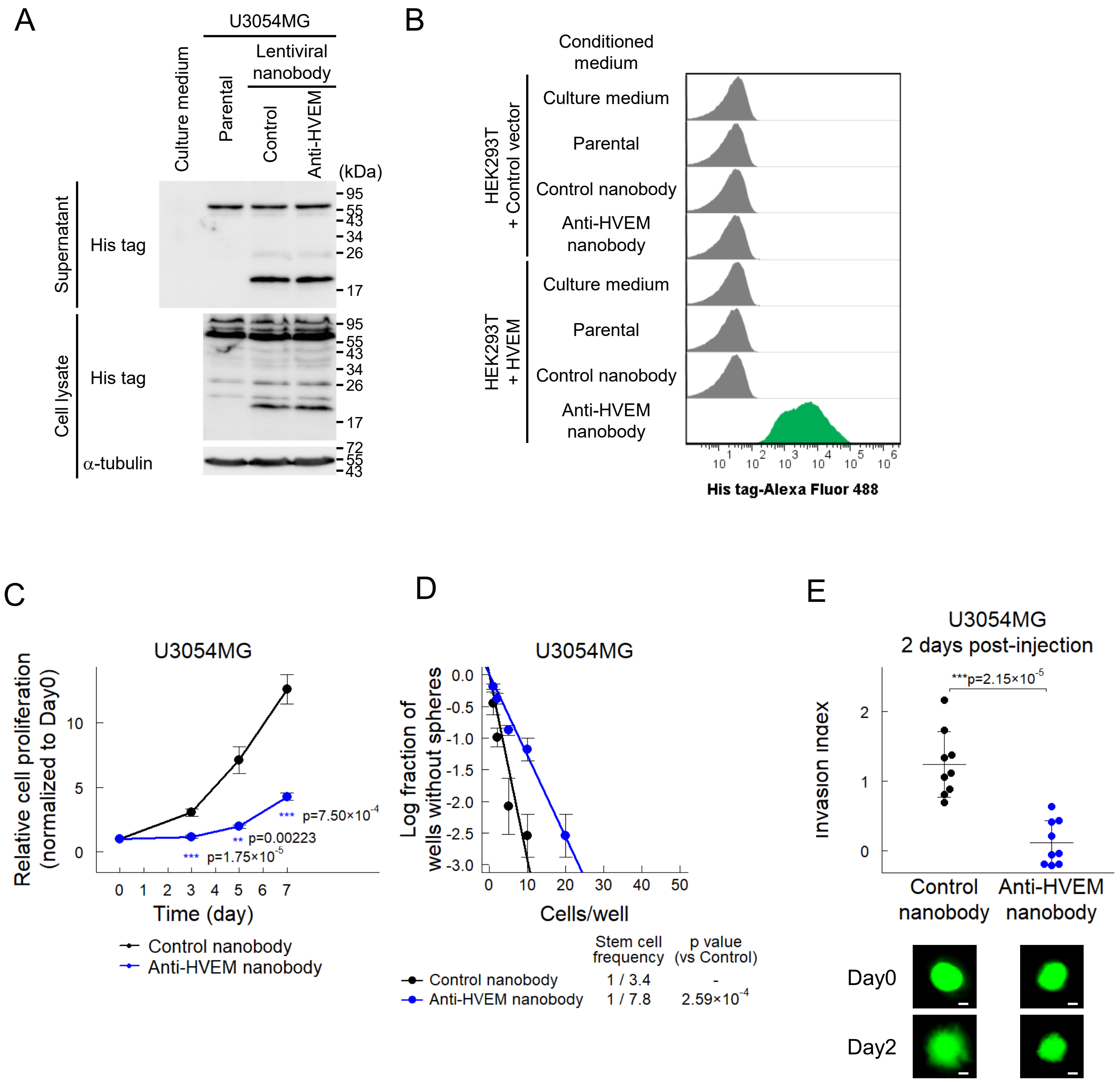

Supplementary Figure S6.

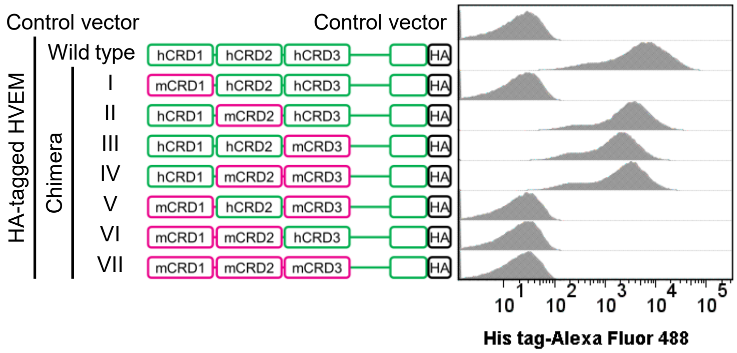

#### SUPPLEMENTARY TABLES

##### Table S1. List of genes incorporated in plasmids (shown in the excel file)

##### Table S2. Nucleotide sequences of shRNAs and sgRNAs

| shRNA | Sequence (sense sequence + stem loop + antisense sequence + terminator) |
|---|---|
| Control shRNA | AUGGUUUACAUGUUGUGUGAacgugugcuguccguUCACACAACAUGUAAACCAUUUUUU |
| TNFRSF14 shRNA #1 | GUGCAGUCCAGGUUAUCGUGUacgugugcuguccguACACGAUAACCUGGACUGCACUUUUU |
| TNFRSF14 shRNA #2 | GAGCUGACGGGCACAGUGUGUacgugugcuguccguACACACUGUGCCCGUCAGCUCUUUUU |
| TNFSF13 shRNA #1 | GAGACUCUAUUCCGAUGUAUAacgugugcuguccguUAUACAUCGGAAUAGAGUCUCUUUUU |
| TNFSF13 shRNA #2 | GGCAAGGGCGAAACUUAACCUacgugugcuguccguAGGUUAAGUUUCGCCCUUGCCUUUUU |

| sgRNA | Sequence (guanine + target sequence + gRNA scaffold + terminator) |
|---|---|
| Control sgRNA | gGUAUUACUGAUAUUGGUGGGguuuuagagcuagaaauagcaaguuaaaauaaggcuaguccguuaucaacuugaaaaaguggcaccgagucggugcUUUUUU |
| TNFRSF14 sgRNA #1 | gAGCAGUUCCGGCUCGCGCGCguuuuagagcuagaaauagcaaguuaaaauaaggcuaguccguuaucaacuugaaaaaguggcaccgagucggugcUUUUUU |
| TNFRSF14 sgRNA #2 | gCCCUCCGGACGUCACCACGGguuuuagagcuagaaauagcaaguuaaaauaaggcuaguccguuaucaacuugaaaaaguggcaccgagucggugcUUUUUU |

##### Table S3. Primer sets for quantitative real-time PCR analysis

| Species | Target genes | Forward / Reverse | Sequence (5’ → 3’) |
|---|---|---|---|
| Homo sapiens | *GAPDH* | Forward | GAAGGTGAAGGTCGGAGTC |
| Homo sapiens | *GAPDH* | Reverse | GAAGATGGTGATGGGATTTC |
| Homo sapiens | *TNFRSF14* | Forward | GTGTCTGCAGTGCCAAATGT |
| Homo sapiens | *TNFRSF14* | Reverse | CCACACACGGCGTTCTCT |
| Homo sapiens | *TNFSF13* | Forward | TATAGCGCAGGTGTCTTCCA |
| Homo sapiens | *TNFSF13* | Reverse | ACAGTTTCACAAACCCCAGGA |
